# Division of labor among H3K4 Methyltransferases Defines Distinct Facets of Homeostatic Plasticity

**DOI:** 10.1101/2023.09.20.558734

**Authors:** Takao Tsukahara, Saini Kethireddy, Katherine Bonefas, Alex Chen, Brendan LM Sutton, Yali Dou, Shigeki Iwase, Michael A. Sutton

**Author notes:** Correspondence (S.I.), (M.A.S.).

## Abstract

Heterozygous mutations in any of the six H3K4 methyltransferases (KMT2s) result in monogenic neurodevelopmental disorders, indicating nonredundant yet poorly understood roles of this enzyme family in neurodevelopment. Recent evidence suggests that histone methyltransferase activity may not be central to KMT2 functions; however, the enzymatic activity is evolutionarily conserved, implicating the presence of selective pressure to maintain the catalytic activity. Here, we show that H3K4 methylation is dynamically regulated during prolonged alteration of neuronal activity. The perturbation of H3K4me by the H3.3K4M mutant blocks synaptic scaling, a form of homeostatic plasticity that buffers the impact of prolonged reductions or increases in network activity. Unexpectedly, we found that the six individual enzymes are all necessary for synaptic scaling and that the roles of KMT2 enzymes segregate into evolutionary-defined subfamilies: KMT2A and KMT2B (fly-Trx homologs) for synaptic downscaling, KMT2C and KMT2D (Trr homologs) for upscaling, and KMT2F and KMT2G (dSet homologs) for both directions. Selective blocking of KMT2A enzymatic activity by a small molecule and targeted disruption of the enzymatic domain both blocked the synaptic downscaling and interfered with the activity-dependent transcriptional program. Furthermore, our study revealed specific phases of synaptic downscaling, i.e., induction and maintenance, in which KMT2A and KMT2B play distinct roles. These results suggest that mammalian brains have co-opted intricate H3K4me installation to achieve stability of the expanding neuronal circuits.

## Introduction

Histone 3 Lysine 4 methylation (H3K4me) is one of the most extensively regulated post-translational modifications in metazoans. The three states, mono-, di-, and tri-methylation (H3K4me1-3), differentially mark gene regulatory elements at transcriptionally-engaged genomic loci. H3K4me3 is enriched at active promoters, whereas H3K4me1 is a hallmark of transcriptional enhancers (Barski *et al*., 2007)(Heintzman *et al*., 2007). A family of enzymes consisting of six distinct lysine methyltransferases -KMT2A, KMT2B, KMT2C, KMT2D, KMT2F, and KMT2G -install H3K4me in mammals(Cenik and Shilatifard, 2021)(Froimchuk, Jang and Ge, 2017)(Rao and Dou, 2015). These six “writer” enzymes appear to have originated from gene duplication of three ancient enzymes found in the fruit fly. KMT2A and KMT2B are fly Trithorax (Trx) homologs, KMT2C and KMT2D are fly Trithorax-related (Trr) homologs, while KMT2F and KMT2G are descendants of *Drosophila* Set (dSet).

Each of the six H3K4me writers are essential for animal development; homozygous deletion of any of the six enzymes in mice leads to embryonic lethality (Bledau *et al*., 2014)(Glaser *et al*., 2006)(Chiu *et al*., 2017)(Lee *et al*., 2013)(Yu *et al*., 1998). The six enzymes likely contribute to animal development by controlling distinct genes and regulatory elements - Trx-related enzymes counteract the polycomb repressive complex to control Hox gene expression during body patterning(Yu *et al*., 1995)(Yu *et al*., 1998); Trr-related enzymes promote gene expression programs for cellular differentiation genes by establishing and maintaining transcriptional enhance(Herz *et al*., 2012)(Hu *et al*., 2013)(Lee *et al*., 2013); and, dSet-related enzymes are responsible for H3K4me3 at the promoters of genes involved in early embryogenesis and blood cell differentiation(Bledau *et al*., 2014)(Tusi *et al*., 2015)(Li *et al*., 2016).

Despite these features of H3K4me writers, it has proven difficult to define a clear role for H3K4 methylation *per se* in nervous system development. In addition to their methyltransferase activity, there is clear evidence that non-enzymatic functions of KMT2 family members are important (Morgan and Shilatifard, 2020). *Kmt2a* null mice are early embryonic lethal, whereas mice lacking only the SET catalytic domain are viable, albeit with some hox gene misregulation (Terranova *et al*., 2006). Trr-null fly mutants are embryonic lethal (Sedkov *et al*., 1999), whereas the mutant complemented with catalytically-inactive Trr is viable with a subtle morphological abnormality (Rickels *et al*., 2017). KMT2C and KMT2D double null mESCs display much more severe transcriptome alteration than mESCs with double KMT2C and KMT2D mutations that only ablate catalytic activities (Dorighi *et al*., 2017)(Wang *et al*., 2016). Independent of catalytic function, KMT2C and KMT2D can recruit histone acetyltransferases and regulate RNA polymerase II. The fruit fly mutants, in which H3K4 is substituted with arginine or alanine, survive with intact developmental gene expression, albeit with a smaller body size (Hödl and Basler, 2012). These results imply that H3K4me may be a minor function of KMT2 family members; however, all six enzymes in mammals, including mice and humans, are active, indicating the presence of selective pressure to maintain the enzymatic function.

Strikingly, when heterozygously mutated in humans, all six KMT2 genes cause monogenic neurodevelopmental disorders, including intellectual disability, autism spectrum disorders, and schizophrenia. Haploinsufficiency of *KMT2A* and *KMT2B* is responsible for Wiedemann-Steiner syndrome (Jones *et al*., 2012) and childhood-onset dystonia-28 (Zech *et al*., 2016), respectively. Heterozygous mutations of KMT2C and KMT2D underlie Kleefstra syndrome-2 (Koemans *et al*., 2017) and Kabuki syndrome-1 (Ng *et al*., 2010), respectively. *KMT2F* loss-of-function mutations have been reported in schizophrenia (Takata *et al*., 2014) and early-onset epilepsy (Yu *et al*., 2019). Finally, KMT2G loss-of-function variants explain several patients with an intellectual developmental disorder with seizures and language delay (Hiraide *et al*., 2018). Thus, the human brain cannot tolerate the heterozygous loss of any of the six KMT2-family enzymes.

Here, we define an essential role for H3K4me in a form of synaptic plasticity – homeostatic synaptic scaling - thought to be key for stabilizing activity in developing neural circuits (Turrigiano *et al*., 1998)(Turrigiano, 2012). Impaired homeostatic plasticity is a pathological hallmark of several neurodevelopmental disorders, including Fragile X syndrome, Tuberous Sclerosis, and Smith-Magenis Syndrome (Bateup *et al*., 2013)(Soden and Chen, 2010)(Garay *et al*., 2020). Synaptic scaling is a transcription-dependent process (Turrigiano *et al*., 1998) and requires several chromatin regulators, including TET3 DNA demethylase (Yu *et al*., 2015), MeCP2 methyl-DNA binder (Zhong, Li and Chang, 2012), EHMT1/2 histone H3K9 methyltransferases (Benevento *et al*., 2016), L3MBTL1 methyl-histone binding factor (Mao *et al*., 2018), and RAI1 nucleosome-binding protein (Garay *et al*., 2020). However, the roles of KMT2 enzymes and H3K4me in homeostatic plasticity remain unknown. We show that H3K4me is dynamically regulated during sustained alterations in neural activity, and that depleting H3K4me impairs excitatory synaptic transmission and homeostatic synaptic scaling in hippocampal neurons. Using a combination of RNAi screening, genetics, and engineered small molecule inhibitors, we further demonstrate unique roles for each of the six KMT2 H3K4 methyltransferases in homeostatic scaling and document a clear division of labor among them that segregates into evolutionarily-defined subfamilies. Finally, our results show that a specific H3K4 writer - KMT2A - acts in a narrowly defined time domain to control transcriptional programs underlying homeostatic downscaling and is critical for the induction, but not maintenance, of synaptic downscaling. Together, our findings demonstrate a clear instructive role for H3K4 methylation in linking persistent changes in neural activity with transcriptional control necessary for adaptive synaptic compensation.

## Results

### Shifts in neuronal activity alter global H3K4 methylation in post-mitotic neurons

Previous studies have shown that H3K4 methylation (H3K4me) is dynamically regulated in response to external stimuli both *in vitro* and *in vivo* (Gupta *et al*., 2010)(Benevento *et al*., 2016). To identify a potential role of KMT2 family enzymes (KMT2s) in neuronal function, we first asked whether H3K4me, the product of KMT2s, changes upon shifts in neuronal activity. We treated DIV14 hippocampal neuronal cultures with the GABA-A receptor antagonist bicuculline (Bic; 40 µM) to increase neuronal activity and quantified the intensity of immunolabeled H3K4me1 and H3K4me3 in MAP2-positive neurons over the time course of 24 h. While the overall expression of H3K4me3 was not significantly altered, we found a robust increase in the expression of H3K4me1 0.5 h after Bic treatment which returned to basal levels within 4 h (Figure 1A and 1B). These changes in H3K4me1 could be a consequence of either *de novo* methylation or demethylation of H3K4me2. In either case, these results indicate that H3K4me is highly dynamic during the early periods of chronic neuronal activation.

**Figure 1.**
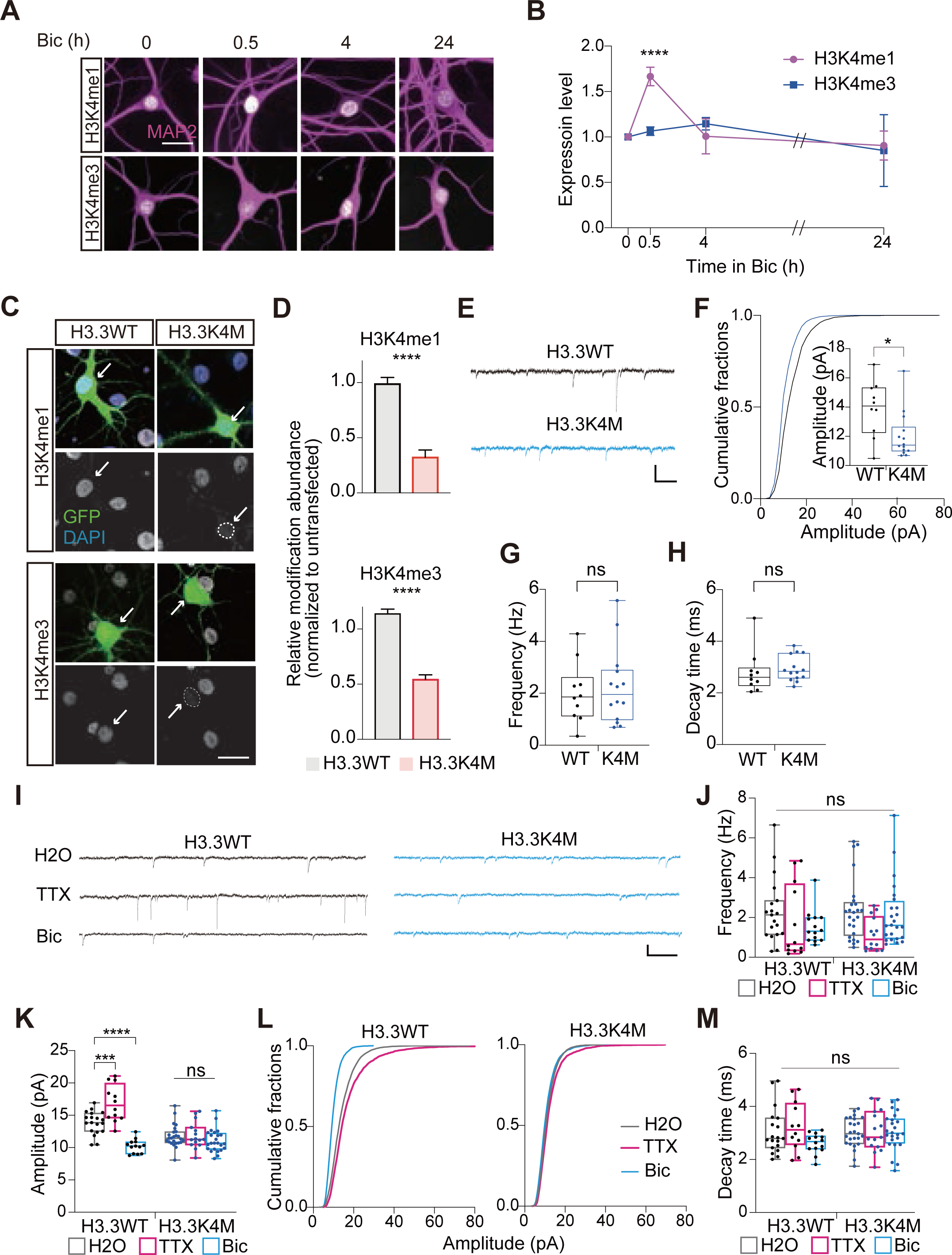
Depletion of H3K4me via ectopic expression of H3.3K4M impairs homeostatic synaptic scaling. (A) Representative images of H3K4me1 (white; top) and H3K4me3 (white; bottom) in MAP2 (magenta) positive neurons upon Bic treatment. Scale bar 20 µm. (B) Line graph of the average intensity of H3K4me1 (magenta) and H3K4me3 (blue), normalized to baseline, in MAP2 positive neurons during Bic treatment. Mean ± S.E.M. ****P < 0.0001. One-way ANOVA followed by post-hoc Tukey’s test. (C) Representative images of H3K4me1 and H3K4me3 in H3.3WT and H3.3K4M transfected neurons (green). White dot lines show H3.3K4M expressing nuclei. Scale bar 20 µm. (D) Bar graph of relative modification abundance of H3K4me1 and H3K4me3 in H3.3WT and H3.3K4M transfected neurons. H3K4me1 (H3.3WT vs. H3.3K4M): N = 20 -17, H3K4me3 (H3.3WT v.s H3.3K4M): N = 57 - 36. Mean ± S.E.M. Unpaired t-test. **** P < 0.0001. (E) Representative miniature excitatory postsynaptic currents (mEPSC) traces from H3.3WT and H3.3K4M. Scale bar 125 ms, 20 pA. (F-H) Box plots of average mEPSC amplitude and the cumulative probability curve of mEPSC amplitude. mEPSC frequency (Hz) and decay time (ms). From here on, box plots indicate the 1st and 3rd quartiles with the median. Whiskers show the minimum and maximum data points. (I) Representative traces of mEPSC from H3.3WT and H3.3K4M transfected neurons treated with H2O, TTX, or Bic for 24∼48 hrs. *P < 0.05, ns: P > 0.05. (J, K, and M) Box plots of average mEPSC frequency, amplitude, and decay time of H3.3WT and H3.3K4M transfected neurons treated with H2O, TTX, or Bic for 24∼48 hrs. H3.3WT: N = 19, 12, 14. H3.3K4M: N = 25, 15, 24 (order: H2O, TTX, Bic). (L) Cumulative probability curve of mEPSC amplitude. One-way ANOVA followed by post-hoc Tukey’s test. ***P < 0.001, ****P < 0.0001. ns: P > 0.05.

### Depletion of H3K4me alters synaptic function and prevents homeostatic synaptic scaling

Since recent studies highlight the importance of non-catalytic functions of KMT2 family members, the causal role of H3K4me at the cellular level remains largely unknown (Rickels *et al*., 2017)(Mishra *et al*., 2014). Therefore, to dissect the role of H3K4me more directly in neurons, we adopted a strategy of depleting this modification in a subpopulation of pyramidal neurons. Lys-to-Met (K-M) mutations have been used for studying the functional impact of site-specific histone methylation events, and a subfraction of these K-M mutants exert a dominant-negative effect to dampen H3Kme globally in multiple cell types (Jang *et al*., 2019)(Herz *et al*., 2014)(Gehre *et al*., 2020). Therefore, to probe the cell-autonomous role of H3K4me, we sparsely transfected either wild-type Histone H3.3 (H3.3 WT), a replication-independent H3 variant, or H3.3 K-M mutant (H3.3K4M) to hippocampal neurons at DIV 2. After 12 d (DIV14), this expression of H3.3K4M reduced H3K4me1 levels by ∼70% and H3K4me3 levels by ∼50% in transfected neurons (H3.3K4M+), compared to untransfected cells. In contrast, H3.3WT transfected neurons (H3.3WT+) exhibited no significant change in H3K4 methylation level compared to untransfected cells (Figures 1E and 1F). This depletion of H3Kme was specific for the H3K4 residue as methylation of neighboring lysine residues -such as H3K9, H3K27, and H3K36 - were unaffected (Figure S2).

With the successful depletion of H3K4me, we tested whether H3K4me has a role in basal glutamatergic transmission. We performed whole-cell voltage-clamp recordings to measure α-amino-3-hydroxy-5-methyl-4-isoxazolepropionic acid receptor (AMPAR)-mediated miniature excitatory post-synaptic currents (mEPSCs), an indicator of excitatory synaptic strength, in H3.3WT+ and H3.3K4M+ pyramidal neurons at DIV 14. We found that H3K4me depletion induced a substantial decrease in mEPSC amplitude in H3.3K4M+ neurons compared to H3.3WT+ neurons, with no detectable difference in mEPSC frequency and decay time (Figure 1 E-H). These results form the first causal role of H3K4me in post-mitotic neurons and suggest potent roles of H3K4me in controlling excitatory synapse function.

The dynamic changes in H3K4me during shifts in neuronal activity led us to test whether H3K4me plays a functional role during synaptic scaling, a transcription-dependent form of homeostatic plasticity (Turrigiano *et al*., 1998)(Turrigiano, 2012). Synaptic scaling is a compensatory mechanism in which neurons alter their synaptic strength to buffer persistent changes in neuronal activity to maintain the stability of firing rates in neural circuits. We treated primary hippocampal cultures with either 1 µM TTX (to block firing) or 40 µM Bic (to persistently enhance firing) for 24 h at DIV 14 to induce synaptic scaling. As expected, TTX and Bic treatment increased and decreased mEPSC amplitude in H3.3WT+ neurons, respectively (Figure 1K), without affecting mEPSC frequency or decay time (Figure 1J, M). The multiplicative shifts in the distribution of mEPSC amplitude in H3.3WT+ control neurons, a hallmark of synaptic scaling, were readily apparent (Figure 1L). Conversely, neither TTX nor Bic treatment changed mEPSC amplitude or shifted the mEPSC cumulative amplitude distribution in H3.3K4M+ neurons (Figure 1K-L). These observations suggest that H3K4me plays a key role in the induction of homeostatic synaptic scaling. Of note, however, depleting H3K4me diminishes basal mEPSC amplitude, so the impact of a floor effect on synaptic downscaling cannot be fully ruled out in this experiment. A role for H3K4me in synaptic downscaling is firmly established by additional experiments targeting individual KMT2 enzymes (see Figures 2-5).

**Figure 2.**
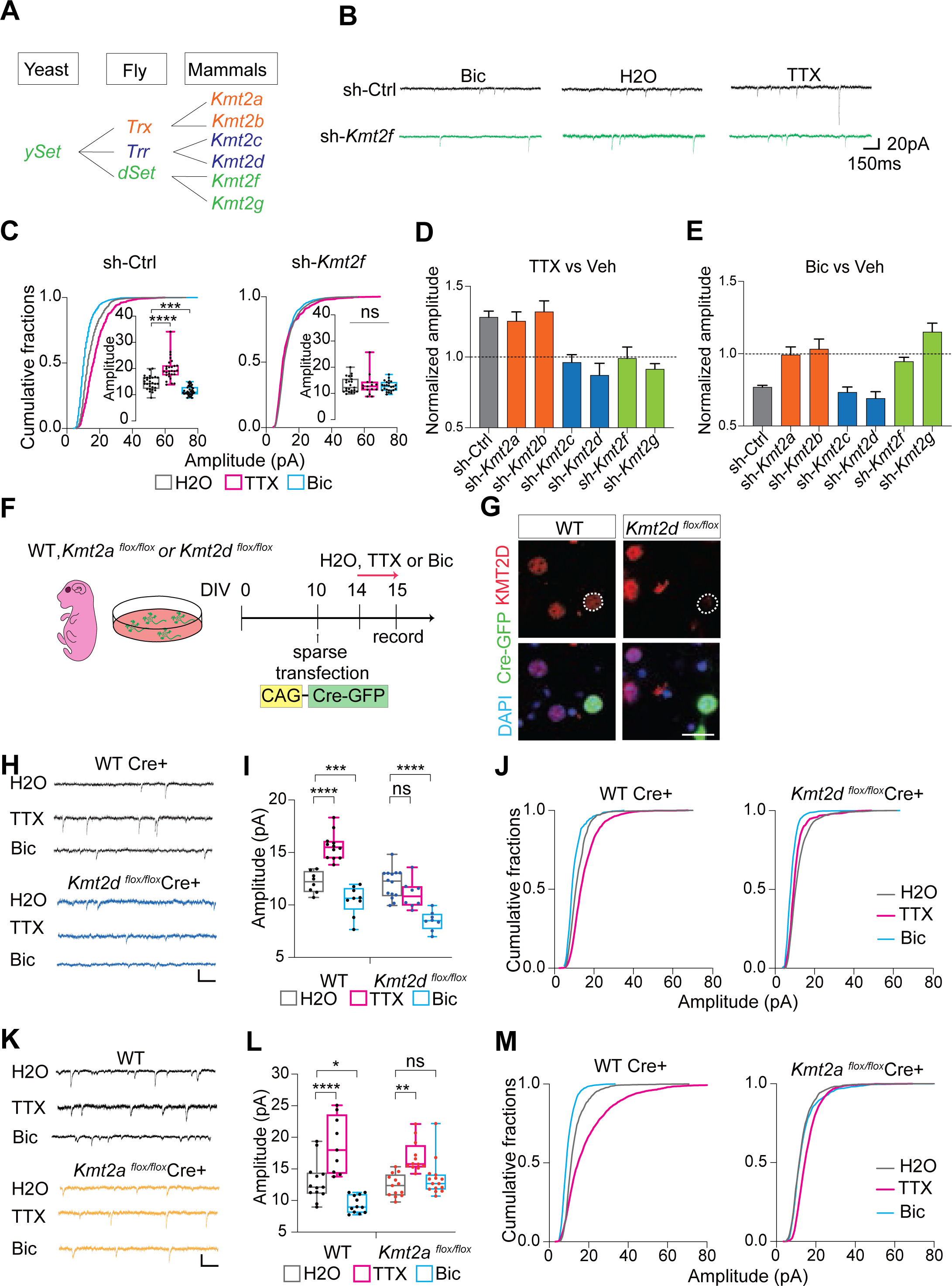
RNA interference screening and conditional knockout of KMT2s reveals a unique and non-redundant role of individual members in synaptic scaling. (A) Evolutional trajectory of H3K4 methyltransferases in Yeast, Fly, and mammals. Fly Trx, Trr, and dSet groups, including their mammalian orthologs, are labeled with orange, blue and green, respectively. (B) Representative mEPSC traces from sh-Ctrl and sh-*Kmt2f* transfected neurons treated with H2O, TTX, or Bic for 24∼48 h. (C) Box plots and cumulative probability curve of mEPSC amplitude for sh-Ctrl and sh-*Kmt2f* transfected neurons treated with H2O, TTX, or Bic for 24∼48 hrs. sh-Ctrl: N = 29, 28, 37. sh-*Kmt2f*: N = 23, 15, 25 (order: H2O, TTX, Bic). One-way ANOVA followed by post hoc Tukey’s test. ***P < 0.001, ****P < 0.0001. ns: P > 0.05. (D and E) Effect of a KD of a single member in the KMT2 family in upscaling and downscaling. Bar graph of average mEPSC amplitude of TTX (D) and Bic (E) groups normalized by the average of H2O groups for neurons treated with a sh-RNA targeting a specific member in the Kmt2s. (D) A value above 1 indicates an upscaling effect. (E) A value below 1 indicates a downscaling effect. sh-Ctrl (gray), Trx-group (*Kmt2a* and *Kmt2b*): orange, Trr-group (sh-*Kmt2c* and *Kmt2d*): blue, dSet-group (sh-Kmt2f and sh-Kmt2g): green. (F) Schematic of experimental procedure. (G) Representative images of KMT2D (red) in neurons derived from WT and *Kmt2d*^flox/flox^ mice. The bottom panel shows merge images of Cre-GFP (green), DAPI (blue), and KMT2D (red). Scale bar 20 µm. (H) Representative traces of mEPSC from WT and *Kmt2d* ^flox/flox^ mice expressing Cre-GFP. Scale bar 20 pA, 150 ms. (I) Box plots of average mEPSC amplitudes from WT (black) and *Kmt2d ^flox^*^/flox^ (blue) mice neurons expressing Cre-GFP. Neural cultures were treated with H2O, TTX, or Bic for 24∼48 hrs. WT: N = 8, 12, 9. *Kmt2d* ^flox/flox^: N = 14, 8, 8 (order: H2O, TTX, Bic). One-way ANOVA followed by post hoc Tukey’s test. ***P < 0.001, ****P < 0.0001. ns: P > 0.05. (J) Cumulative probability curve of mEPSC amplitude from WT and *Kmt2d^flox^*^/flox^ mice neurons expressing Cre. (K) Representative traces of mEPSC from WT (black) and *Kmt2a^flox^*^/flox^ (orange) mice expressing Cre-GFP. Scale bar 20 pA, 150 ms. (L) Box plots of average mEPSC amplitudes from WT and *Kmt2a^flox^*^/flox^ mice neurons expressing Cre-GFP. Neural cultures were treated with H2O, TTX, or Bic for 24∼48 hrs. WT: N = 13, 12, 9. *Kmt2a^flox^*^/flox^: N = 13, 15, 12 (order: H2O, TTX, Bic). One-way ANOVA followed by post hoc Tukey’s test. *P < 0.05, **P < 0.01, ****P < 0.0001. ns: P > 0.05. (M) Cumulative probability curve of mEPSC amplitude from WT and *Kmt2a^flox^*^/flox^ mice neurons expressing Cre-GFP.

### An RNAi screen reveals a division-of-labor among KMT2 H3K4 methyltransferases in synaptic scaling

Given that H3K4 methylation is required for synaptic scaling, we sought to ask which member of the KMT2 family plays a role in this plasticity. First, using publicly available single-cell RNA-seq datasets, we confirmed that all six genes in the KMT2 family (KMT2s) are present in the majority of excitatory neurons in the hippocampus (Figure S1). Next, we employed an RNA interference (RNAi)-screening strategy to acutely knockdown (KD) individual *Kmt2s* in a subpopulation of pyramidal neurons. This approach allowed us to dissect the cell-autonomous roles of each enzyme individually without the impact of gene manipulation in synapse development. Therefore, we transfected neurons with either non-targeting scramble shRNA or shRNAs specific for each *Kmt2* family member at DIV 12. At DIV 14, neurons were treated with either TTX or Bic for 24 h to induce synaptic scaling. KD of individual genes in *Kmt2s* revealed a modest effect, if any, on basal mEPSC features relative to scramble shRNA treated neurons (Figure S3).

As expected, neurons transfected with scrambled shRNA showed robust up- and down-scaling reflected by increases and decreases in the amplitude of mEPSCs in response to TTX and Bic, respectively (Figure 2B). We anticipated that KD of most KMT2 family members individually would have minimal effects on up- and down-scaling, given the potential functional redundancy conferred by the intact expression of the other five family members. Surprisingly, however, we found that KD of each KMT2 enzyme impacted synaptic scaling and that specific phenotypes associated with loss of each KMT2 emerged. For example, KMT2A- and KMT2B-depleted neurons exhibited robust upscaling in response to TTX, but failed to downscale mEPSC amplitudes following Bic treatment (Figure 2B-C). By contrast, KMT2C and KMT2D KD neurons showed normal downscaling of mEPSC amplitude in response to Bic, but no upscaling of mEPSC ampltidue following TTX treatment (Figure 2B-C). Finally, KD of KMT2F and KMT2G prevented scaling of mEPSC amplitude in both directions, implying their critical role in both up- and -down-scaling (Figure 2B-C). These results demonstrate that all six KMT2 writers play non-redundant roles in the two modes of synaptic scaling. When considering the evolution of the KMT2 family, an intriguing pattern emerges – all six KMT2 family members originate from *ySet* in yeast, the sole H3K4 methyltransferase in the organism, but diverge in multi-cellular organisms such as *Drosophila*. KMT2A and KMT2B share *Drosophila* Trx as their ancestral protein, while KMT2C and KMT2D originate from Trr, and the remaining KMT2F and KMT2G are structurally related to dSet (Figure 2A). Thus, the labor between KMT2 H3K4 methyltransferases in synaptic scaling segregates into evolutionally defined subfamilies.

### Conditional deletion of *Kmt2a* and *Kmt2d* impairs homeostatic down-scaling and up-scaling, respectively

To validate our findings based on RNAi screening, we used a conditional genetics approach. We asked whether conditional deletion of individual KMT2 family members, again in a context where the remaining KMT2s remain intact, would recapitulate the specific phenotypes we observed in homeostatic synaptic scaling. We made primary hippocampal cultures from two flox lines - *Kmt2a ^flox^* and *Kmt2d ^flox^*, representing *Trx* and *Trr* groups, respectively –and their WT littermates. Both WT and floxed neurons were transfected with Cre-GFP at DIV10, and used for experiments at DIV14. This time period allowed for robust depletion of the target genes in the floxed alleles, as revealed by immunocytochemistry comparing transfected and untransfected *Kmt2d ^flox/flox^*neurons at DIV 14 (Figure 2H). Using this conditional genetic strategy, we asked whether deletion of KMT2A and KMT2D would recapitulate the same scaling phenotypes evident after RNAi-mediated knockdown. As expected, we found that Cre-GFP+ WT neurons exhibited robust up- and downscaling in response to 24-48 h treatment with either 1 µM TTX or 40 µM Bic, respectively (Figure 2I-K). Similar to our RNAi findings, we found that conditional deletion of KMT2D in Cre-GFP+ neurons blocked upscaling, but not downscaling (Figure 2I-K), while deletion of KMT2A produced the opposite pattern of results (Figure 2L-N). Recordings from neighboring untransfected *Kmt2d ^flo/flox^* or *Kmt2a ^flox/flox^* neurons revealed intact up- and downscaling, respectively in the absence of Cre-mediated recombination (Figure S3H-K). These results confirm a division of labor among H3K4me writers in instantiating compensatory synaptic adaptations in response to chronic alterations in network activity.

### KMT2A Regulates Synaptic Down-scaling via the Catalytic SET Domain

As noted, knockout of individual KMT2 family members leads to embryonic lethality, but knock-in animals expressing a methyltransferase-dead KMT2 at the native locus survive and have relatively mild phenotypes. This pattern implies that in most cell types, the methyltransferase activity among KMT2 family members has substantial functional redundancy but the non-enzymatic functions are unique. However, the enzymatic SET domain in all six enzymes is conserved through evolution suggesting that there has been strong selection pressure to maintain the enzymatic activity of each KMT2 member. Of relevance, single allele mutations in each KMT2 family member are associated with neurodevelopmental disorders, suggesting that this selection pressure may relate to important and unique functional roles of neuronal methylation conferred by each specific KMT2. If so, then removing just the methyltransferase activity of specific KMT2 enzymes should produce similar synaptic effects as removing the whole protein. To examine this possibility, we employed a knock-in KMT2A mouse model harboring a deletion of the catalytic SET domain (*Kmt2a*ΔSET; Figure 3A). We made primary hippocampal cultures from *Kmt2a*ΔSET and their WT littermates and examined synaptic function at DIV 14. We first evaluated basal synaptic transmission and found no detectable change in mEPSC amplitude, frequency, or decay time between the genotypes (Figure S4C-F). By contrast, while WT neurons exhibited increased and decreased mEPSC amplitude in response to TTX and Bic, respectively, *Kmt2a*ΔSET neurons demonstrated only upscaling in response to TTX treatment without any downscaling of mEPSC amplitude after Bic treatment (Figure 3B-D). We found no significant changes in mEPSC frequency in any experimental condition (Figure 3E). Since SET domain mutations can potentially destabilize the protein (Jang et al., 2019), we quantified the expression of KMT2A in neurons. Surprisingly, we observed a mild (but significant) increase in KMT2A expression in Kmt2aΔSET vs WT neurons (Figure S4A-B). While the mechanism underlying this altered expression remains unknown, the results clearly rule out a loss of KMT2A expression as an underlying contributor to the synaptic phenotype. Hence, unlike the functional redundancy observed in other cell types, the methyltransferase activity of KMT2A in neurons appears to play a unique role in homeostatic downscaling that other KMT2 family members cannot compensate for.

**Figure 3.**
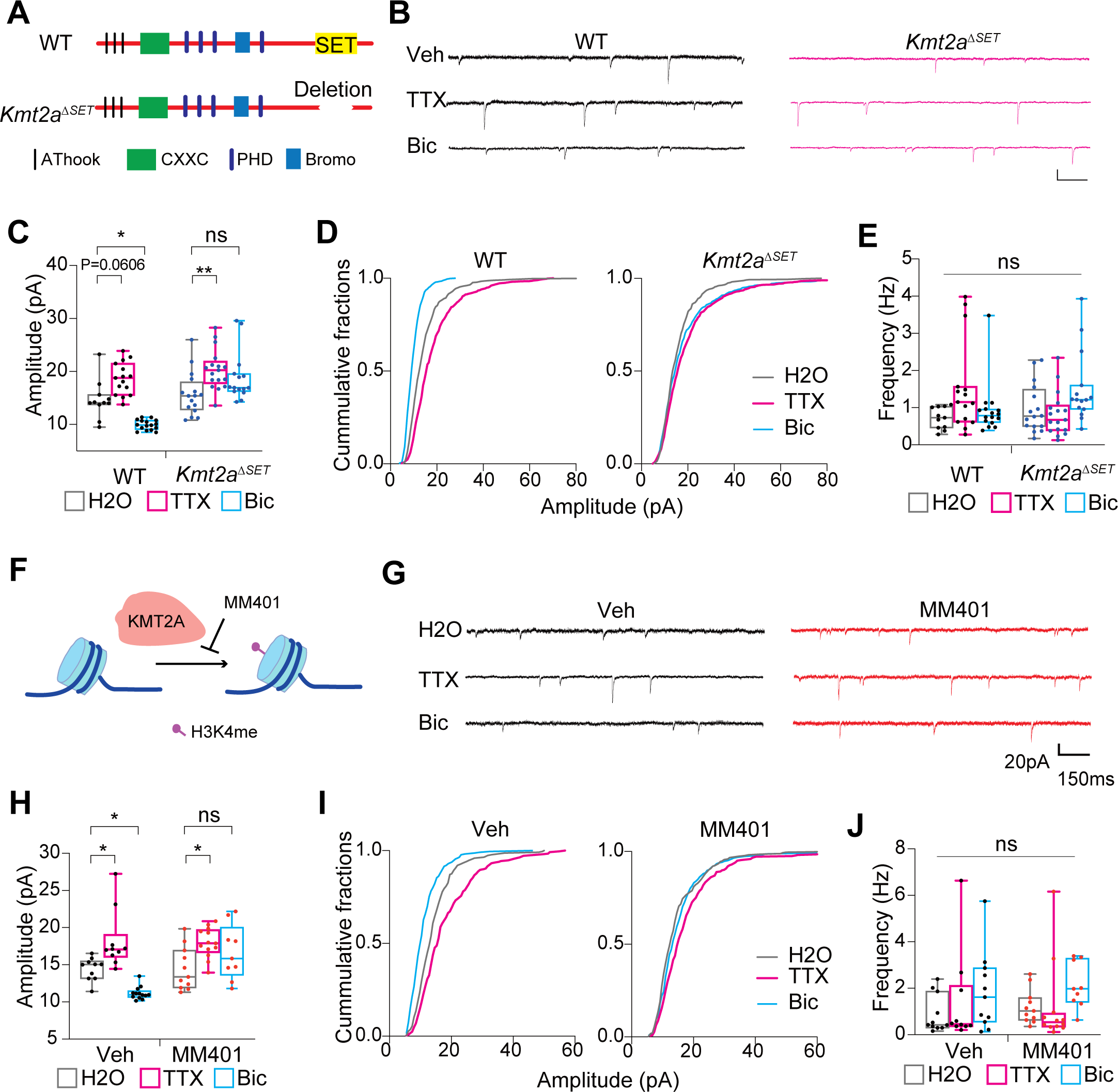
KMT2A mediated activity-dependent H3K4 methylation is required for downscaling. (A) Cartoon of the structure of catalytic dead *Kmt2a* (*Kmt2a*ΔSET) mouse line. (B) Representative traces of mEPSC from WT (black) and Km2a *Kmt2a*ΔSET (pink) mice treated with H2O, TTX, or Bic for 24∼48 hrs. Scale bar 20 pA, 150 ms. (C and E) Box plots of average mEPSC amplitudes(C) and frequency (E) from WT and Kmt2a delta-set mice neurons. Neural cultures were treated with H2O, TTX, or Bic for 24∼48 hrs. WT: N = 11, 15, 16. *Kmt2a*ΔSET: N = 16, 17, 15 (order: H2O, TTX, Bic). One-way ANOVA followed by post hoc Tukey’s test. *P < 0.05, **P < 0.01. ns: P > 0.05. (D) Cumulative probability curve of mEPSC amplitude from WT and *Kmt2a*ΔSET mice neurons. (F) Cartoon of MM401 inhibiting the catalytic function of KMT2A. (G) Representative mEPSC traces of vehicle(black) and MM401(red) treated neurons with H2O, TTX, or Bic for 24∼48 hrs. Scale bar 20 pA, 150 ms. (H) Box plots of average mEPSC amplitudes(C) and frequency (E) from WT and *Kmt2a*ΔSET mice neurons. Neural cultures were treated with H2O, TTX, or Bic for 24∼48 hrs. Vehicle: N = 10, 10, 13. MM401: N = 11, 13, 9 (order: H2O, TTX, Bic). One-way ANOVA followed by post hoc Tukey’s test. *P < 0.05. ns: P > 0.05. (I) Cumulative probability curve of mEPSC amplitude from vehicle and MM401 treated neurons.

### Activity-dependent H3K4 methylation is required for synaptic downscaling

Both acute knockdown and genetic experiments establish a specific role for KMT2A in homeostatic downscaling, raising two possible scenarios: 1) chronic loss of H3K4me installed by KMT2A creates a non-permissive state for inducing homeostatic downscaling or 2) the acute activity-dependent deposition of H3K4me by KMT2A plays an instructive role. To distinguish between these possibilities, we took advantage of a small molecule inhibitor - MM401 - recently developed by our group, which specifically inhibits the enzymatic activity of KMT2A without affecting the enzymatic activity of other KMT2 family members (Figure 3F; Cao et al., 2014). We pretreated rat hippocampal neurons (DIV14) with 25 µM MM401 or vehicle 0.5 h prior to and throughout 24 h treatment with either Bic or TTX. Consistent with our earlier studies, we found that acute KMT2A inhibition with MM401 completely blocked down-scaling induced by Bic, but not up-scaling following TTX treatment (Figure 3G-I). We did not observe any effect on basal mEPSC amplitude in the absence of activity shifts (Figure 3H), nor did we find significant changes in mEPSC frequency with MM401 treatment (Figure 3J). Together, these results demonstrate that the acute activity-dependent installation of H3K4me plays an instructive role in homeostatic synaptic downscaling.

### KMT2A is required for the induction, but not maintenance, of synaptic downscaling

MM401 is a tool that allows to acutely disrupt KMT2A activity, giving us the opportunity to probe how KMT2A-driven H3K4me contributes to the transcriptional program underlying homeostatic downscaling. While chronic changes in neural activity are required to induce homeostatic scaling, the resulting synaptic adaptations are presumably maintained in the face of continued conditions that would otherwise lead to hyperactivation of the network (Figure 4A). However, very few studies have distinguished mechanisms that operate in the induction vs maintenance phase of homeostatic plasticity. This distinction is likely important as a recent study demonstrated that the inability to maintain presynaptic homeostatic plasticity, another form of homeostatic plasticity, leads to disease progression in an amyotrophic lateral sclerosis model (Orr *et al*., 2020).

**Figure 4.**
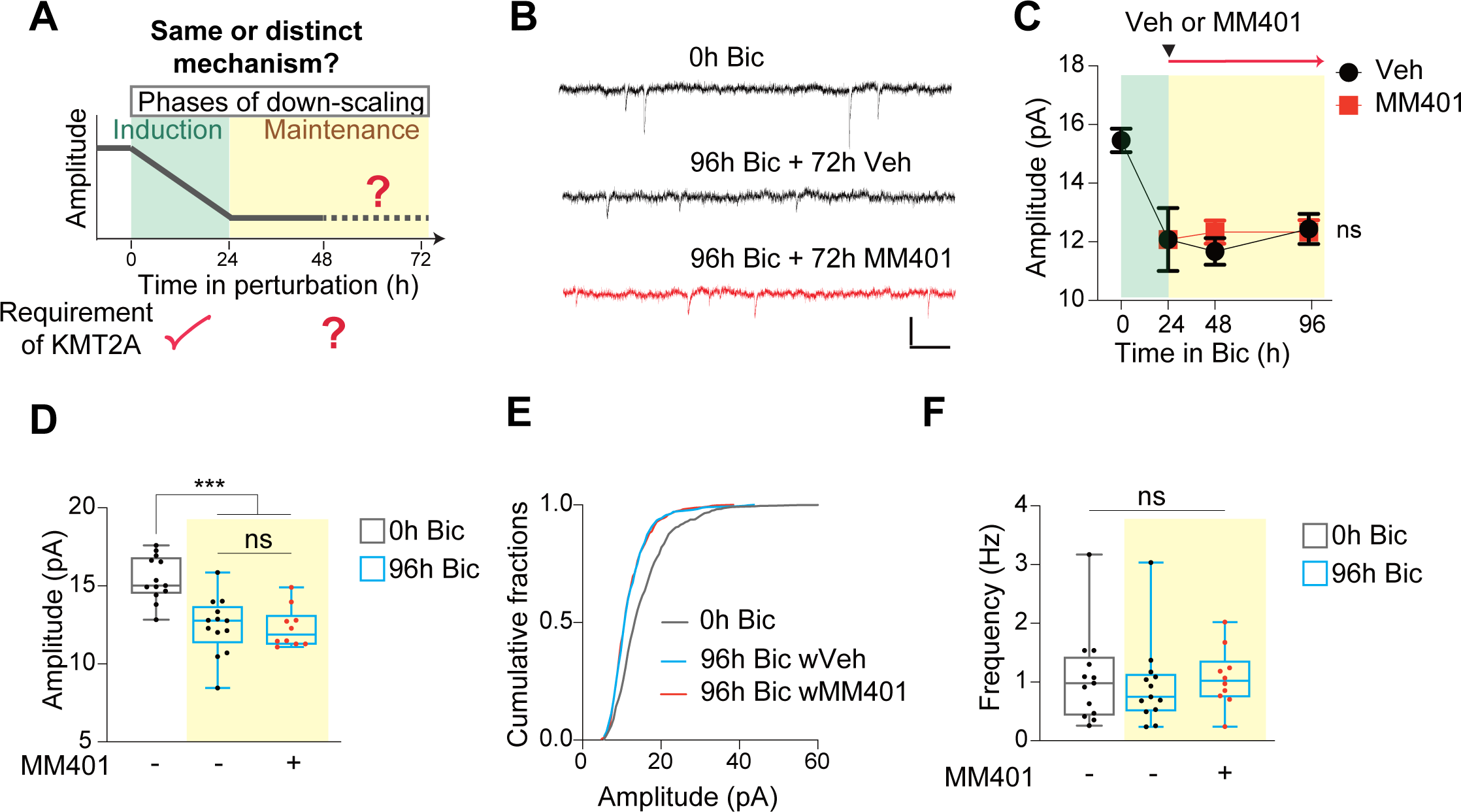
Maintenance of homeostatic downscaling does not require KMT2A. (A) Concept of induction and maintenance phase of synaptic downscaling. The induction of downscaling is established within 24 h after perturbation. This induction requires enzymatic activity of KMT2A, however, whether and how downscaling is maintained remains largely unknown. (B) Representative traces of mEPSC with 0 h, 96 h Bic with or without 72 h vehicle, or MM401. Scale bar 20 pA, 150 ms. (C) Time course of mEPSC amplitude during synaptic downscaling. Black and red lines are vehicle and MM401 treated groups. Mean ± S.E.M. ns > 0.05. Two-way ANOVA followed by post hoc Bonforreni’s test. (D and E) Box plot and cumulative probability curve of mEPSC amplitude comparing 0 h and 96 h Bic with or without 72 h MM401. One-way ANOVA followed by Tukey’s multiple comparisons tests. ***P < 0.001. ns: P > 0.05. (F) Box plot of average mEPSC frequency comparing 0 h Bic and 96 h Bic with or without MM401. One-way ANOVA followed by Tukey’s multiple comparisons tests. ns: P > 0.05.

Using MM401, we asked whether KMT2A-driven H3K4me is required specifically for the induction of synaptic downscaling or whether downscaling is continuously dependent on KMT2A for synaptic adaptations to persist. We induced synaptic down-scaling via 24 h Bic treatment in rat hippocampal cultures and subsequently treated neurons with either vehicle or 25 µM MM401 while continuing the Bic treatment for a total of 96 h. mEPSCs were recorded at multiple time-points. As expected, we observed robust downscaling of mEPSC amplitude at 24 h Bic treatment, and this effect persisted for the next 72 h in vehicle-treated neurons (Figure 4B-C, 5SA-C). As with downscaling evident 24 h following Bic, the cumulative probability distribution of mEPSCs after 96 h Bic exhibited a multiplicative shift to the left, a hallmark of synaptic downscaling. We observed a near identical profile of mEPSC amplitude changes in neurons treated with MM401 during the maintenance phase of downscaling (Figure 4B-E), despite the fact that MM401 administration prior to Bic treatment completely blocks the emergence of downscaling (Figure 3G-I). Neither Bic treatment nor MM401 administration produced any significant changes in mEPSC frequency and decay time (Figure 4B-C and 5SA-E). These results establish the existence of a maintenance phase in synaptic down-scaling that is mechanistically distinct from the processes that induce synaptic adaptations. Acute deposition of H3K4me by KMT2A is one important mechanism that links chronic changes in neural activity with the gene expression changes necessary to induce adaptive synaptic weakening. Once these changes are instantiated, however, KMT2A is no longer required to maintain the synaptic changes.

### KMT2A acts in a specific time window for the induction of synaptic downscaling

Having established that KMT2A is selectively required for the induction, but not maintenance, of synaptic downscaling, we next sought to dissect the time window in which KMT2A is acting. First, we asked whether inhibition of KMT2A affects global H3K4 methylation either on its own or in response to an increase in neuronal activity. To this end, we pretreated neurons with either vehicle or 25 µM MM401 0.5 h before Bic treatment and evaluated the intensity of immunolabeled H3K4me1. As before, we observed a robust increase in H3K4me1 in vehicle-treated neurons 0.5 h following hyperactivation, and this Bic-induced increase in H3K4me1 was completely blocked by MM401 pre-treatment; MM401 had no effect on H3K4me1 levels in the absence of network hyperactivation (Figure 5A-B). In addition to this result, we confirmed that Kmt2aΔSET mutant neurons also failed to elevate H3K4me1 levels upon increased neuronal activity (Figure S5F and G). These results suggest that KMT2A is recruited early during activity elevation and is selectively responsible for the resulting early changes in H3K4me.

**Figure 5.**
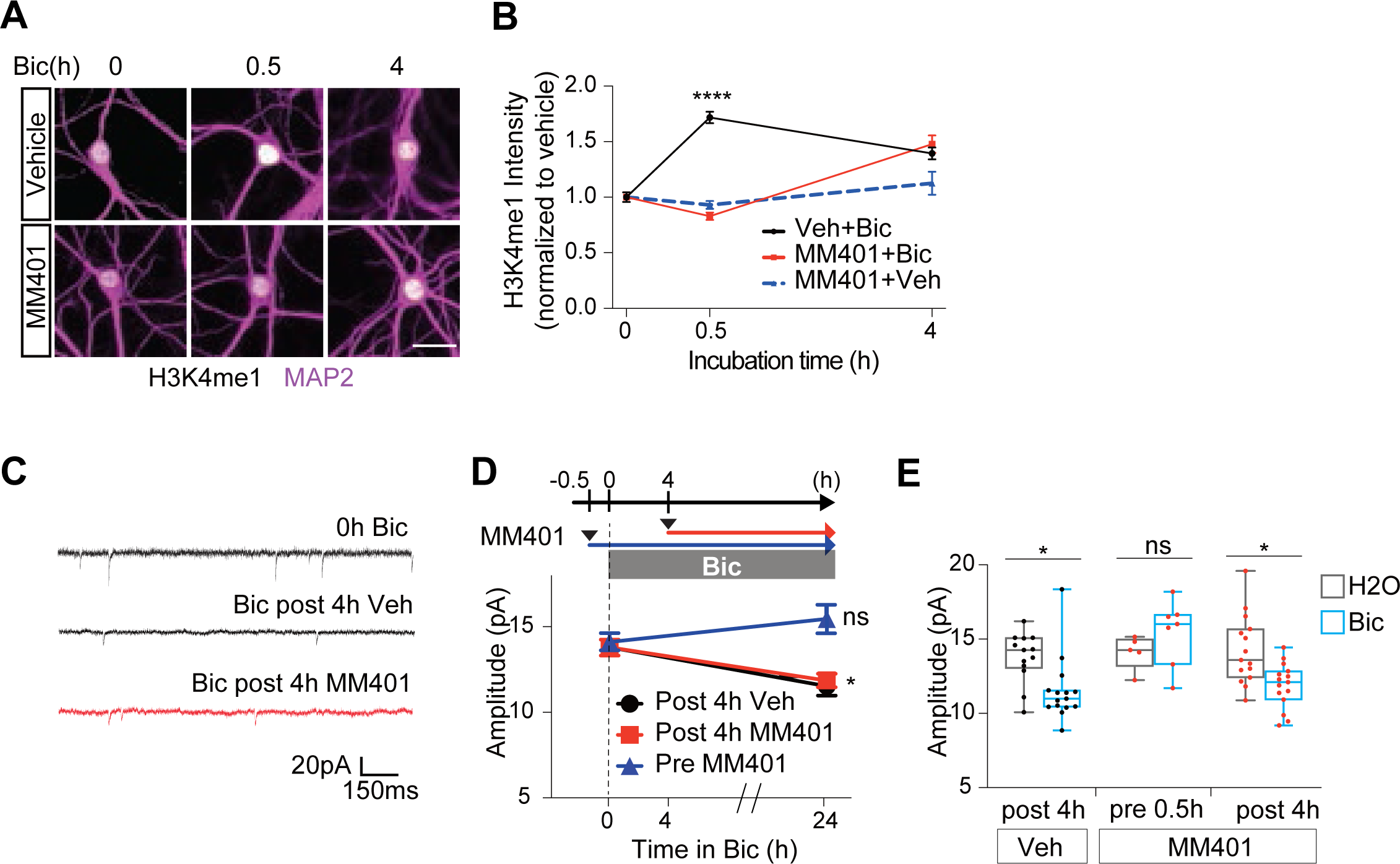
KMT2A is temporally required for homeostatic downscaling. (A) Representative images showing the time course effect of MM401 on H3K4me1 (white; upper) in MAP2 (magenta) positive neurons upon Bic treatment. Scale bar 20 µm. (B) Time course of the average intensity of H3K4me1 in MAP2 positive neurons during Bic treatment. Mean ± SEM. ****P < 0.0001. One-way ANOVA followed by post hoc Tukey’s test. (C) Representative traces of mEPSC with 0 h Bic, 24 h Bic followed by an additional treatment of vehicle or MM401 after 4 h Bic treatment. Scale bar 20 pA, 150 ms. MM401 after 4h Bic no longer impairs downscaling. (D) Schematic of the experimental procedure with the line graph of average mEPSC amplitude. 24 h Bic with vehicle or MM401 treatment after 4 h Bic both show robust downscaling. On the other hand, 0.5 h pretreatment of MM401 before Bic completely blocked the induction of downscaling. Mean ± SEM. *P < 0.05. ns: P > 0.05. One-way ANOVA followed by post hoc Tukey’s multiple comparison test. (E) Box plot of average mEPSC amplitude in panel D.

H3K4me dynamics following activity elevation raises the possibility that H3K4me enzymes are recruited in stages following activity shifts, which would suggest that the requirement for specific enzymes such as KMT2A would be limited to specific time domains. To address this question, we treated hippocampal cultures with vehicle or 25 µM MM401 either prior to (0.5 h before) or 4 h following Bic administration. 24 h following Bic treatment, we observed robust downscaling of mEPSC amplitude in vehicle-treated neurons; this downscaling was nearly identical in neurons treated with MM401 4 h post-Bic, but was completely suppressed in neurons treated with MM401 prior to Bic (Figure 5C-E). These observations establish an important, but transient, role for KMT2A in the induction of synaptic downscaling, with KMT2A only required over the first 4 h of hyperactivity. Given the strong association between H3K4me and transcriptional control, these findings suggest that KMT2A serves as an early link between chronic changes in neural activity and altered gene expression necessary for adaptive changes in synaptic function.

### Inhibition of KMT2A activity impairs the transcriptional program underlying downscaling

Synaptic downscaling relies on de novo transcription induced by an increase of the network activity (Horvath *et al*., 2021). We sought to determine the contribution of KMT2A in the transcriptional program responsible for synaptic scaling. To monitor *bona fide* transcriptional dynamics, instead of steady-state RNA, we employed bromouridine sequencing (BrU-seq), a nascent RNA sequencing technique (Paulsen *et al*., 2013) (Paulsen *et al*., 2014). Neural cultures (DIV 17) were treated with either DMSO or MM401, then activity-dependent transcriptions were elicited by Bic treatment. We then labeled nascent RNA for 30-min prior to harvesting (1h and 4h), enriched BrU-containing RNA, and sequenced them (Figure 6A).

**Figure 6.**
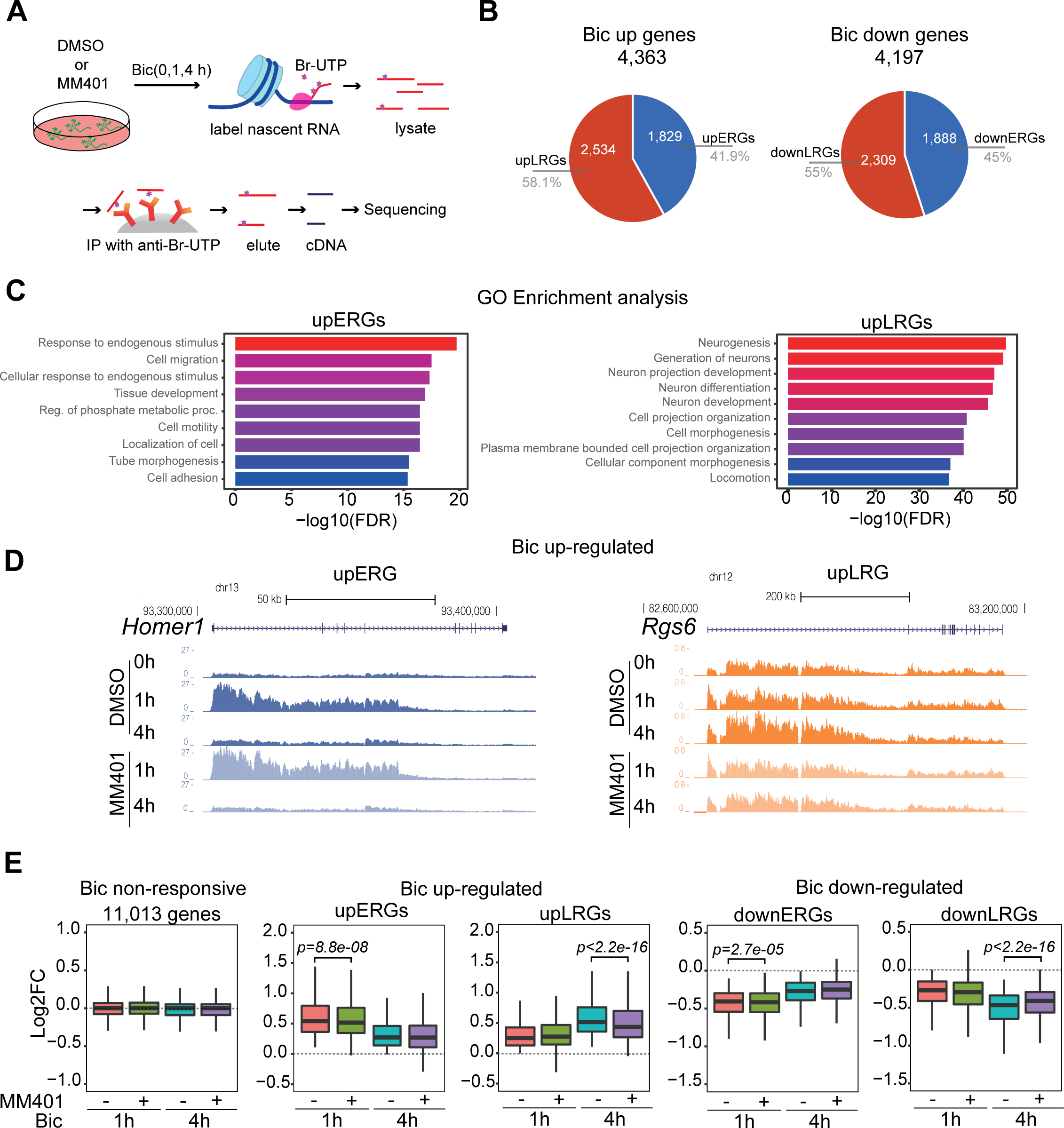
Nascent RNA-seq revealed that KMT2A inhibition attenuates the expression of late-responsive genes. (A) Schematic of BrU-seq. Cortical cultures were treated with either DMSO or 25 µM MM401. Both conditions were followed by Bic or vehicle treatment. All conditions were incubated with bromouridine (brU) for 30 min before lysate to label nascent RNA. Lysates were immune-precipitated with BrU antibody, eluted, reversed transcript to cDNA, and sent to sequencing. (B) Pie-chart of genes categorized by the response to Bic. Bic up-regulated (left) and down-regulated (right) genes were further divided into early-responsive genes (ERGs) and late-responsive genes (LRGs). (C) Gene-ontology enrichment analysis for upregulated ERGs (upERGs) and LRGs (upLRGs). (D) USCS genome browser shot of *Homer1* and *Rgs6*, which represent up-regulated ERG (upERG) and LRG (upLRG), respectively. The expression of *Homer1* is transiently increased upon Bic treatment in DMSO and MM401-treated groups. On the other hand, *Rgs6* expression increased only after 4 h Bic treatment, and this increase was attenuated in MM401-treated groups. The browser shots are from one replicate; however, genes which at least two replicates followed the same trend were selected. (E) Box plot of log2 foldchange upon Bic treatment with or without MM401. Bic non-responsive genes doesn’t have noticeable effect by MM401. The expression of upERGs, upLRGs, downERGs and downLRGs are all attenuated by MM401.

Two waves of gene expression characterize activity-dependent transcriptional programs. The first wave comprises early-response genes (ERGs), expressed rapidly upon neuronal activation, and late-response genes (LRGs), which are slowly expressed and require de novo translation for induction (Fowler, Sen and Roy, 2011). *Arc* and *Bdnf*, well-characterized ERG and LRG, showed an expected increase in transcription evidenced by abundant intronic reads, indicating that Bru-seq captures transcription dynamics reliably (Figure S6A). In our Bru-seq dataset, 4,363 genes were upregulated while 4,197 genes were down-regulated upon either 1hr or 4hr Bic treatment (DEseq2, Padj < 0.05). Of these, 1,829 genes were upregulated in 1hr Bic treatment (upERGs), while 2,534 genes were induced only after 4hr (upLRGs). Leveraging the sensitive detection of transcriptional downregulation by Bru-seq (Garay *et al*., 2020), we also identified transcriptional suppression of 1,888 genes at 1hr (downERGs) and 2,309 genes (downLRGs) only after 4 hr treatment (Figure 6B). While upERGs were enriched with stimuli-dependent ontology terms such as “response to endogenous stimulus”, upLRGs were characterized by neurogenesis and development-related processes. Interestingly, downERGs and downLRGs are strongly enriched with translational regulation, such as RNA processing and ribosome biogenesis, suggesting that activity-dependent transcription is coupled with translational suppression (Figure S6A).

Differential gene expression analysis revealed that the impact of MM401 on transcription is larger at 4 h compared to 1 h Bic treatment; at 1 h post-Bic, 19 differentially expressed genes (DEG) were identified, whereas 240 DEGs are found at 4 h post-Bic (Padj < 0.3, Table S1). Of these DEGs, MM401 impacted 6 ERGs at 1hr compared to 159 LRGs at 4hr (Table S1). A more pronounced impact on LRGs compared to ERGs is exemplified at *Homer1* and *Rgs6* loci, respectively (Figure 6D). We then tested whether MM401 deregulates ERGs and LRGs as groups. MM401 attenuated induction of both upERGs and upLRGs (Wilcoxon signed rank test: p = 8.8e-08 for upERGs and p < 2.2e-16 for upLRGs; Figure 6E). However, the impact of MM401 was substantially larger on LRGs than on ERGs (effect size: d = 2.11 vs. d = 9.55). Interestingly, downERGs and downLRGs were also attenuated by MM401 treatment (Wilcoxon signed rank test: p = 2.7e-05 and p < 2.2e-16; Figure 6E), with a more pronounced effect on the latter (effect size: d = 0.22 vs. d = 11.46). In contrast, Bic-insensitive genes did not show noticeable change upon MM401 treatment at any time point (Figure 6E). These results demonstrate that KMT2A shapes transcriptional responses to increased network activity, especially in the late phase.

## Discussion

Using a combination of genetics, RNAi screening, small molecule inhibitors, and nascent mRNA profiling, this work defines unique roles for the six H3K4 methyltransferases in homeostatic synaptic scaling. As these six enzymes all target a single histone mark, are ubiquitously expressed, and are all related to a single ancestral form found in yeast, the possibility of extensive functional redundancy has made it challenging to understand the specific roles of individual H3K4 writers in brain development. In fact, some recent studies have even called into question the importance of H3K4 methylation to the role of KMT2 family members in development (Terranova *et al*., 2006)(Rickels *et al*., 2017)(Morgan and Shilatifard, 2020), as these enzymes also clearly have important non-enzymatic functions. On the other hand, there has been strong selection pressure to preserve the catalytic activity of each KMT2 family member, suggesting that preserving their unique activity at H3K4 has important functional significance. Our results suggest that, while there may be substantial functional redundancy of KMT2 writers in many tissues, the maintenance of unique H3K4 methyltransferases in neurons is critical for fine-tuning transcriptional outputs to specific patterns of neural activity.

### H3K4 writer’s enzymatic vs. non-enzymatic functions of KMT2 family members

Homozygous loss of any member of the KMT2 family of H3K4 writers is embryonic lethal, yet knock-in animals that express catalytic dead KMT2’s from the native locus survive and, in many tissues, have relatively mild phenotypes. These observations clearly reveal important non-catalytic functions of KMT2 family members, which has made it difficult to resolve functional roles for their methyltransferase activity specifically. One possibility is that the unique writer activity of any one of the six KMT2 methyltransferases is dispensable in many tissues, but critical in others such as the brain. Consistent with this idea, heterozygous mutations in any of the KMT2 family members are associated with neurodevelopmental disorders, suggesting a particularly important neuronal role for each enzyme that the remaining five cannot substitute for. When we depleted H3K4me in hippocampal neurons, via expression of a H3.3K4M mutant, we found robust decreases in basal synaptic strength, as well as a complete loss of homeostatic scaling induced by chronic changes in neuronal activity. Interestingly, basal synaptic strength was mostly unaffected upon individual knockdown of each KMT2 family member, suggesting enough functional redundancy among enzymes to maintain basal synaptic strength at normal levels. On the other hand, homeostatic scaling was consistently disrupted by knocking down (or genetically deleting) individual KMT2 family members, indicating that the unique writer activity of each enzymes plays non-overlapping roles in driving these synaptic adaptations. Importantly, this division of labor among KMT2 enzymes segregated into evolutionarily-defined sub-families – with the Trx family (KMT2A-B) playing specific roles in downscaling, the Trr family (KMT2C-D) with specific roles in upscaling, and the dSET family (KMT2F-G) required for both up- and downscaling. Given the effects of depleting H3K4me, the similar phenotype in *Kmt2a*ΔSET neurons, and the prevention of downscaling by acute enzymatic inhibition of KMT2A, this division of labor appears tied to the unique effects of each enzyme on H3K4me.

### Time-dependent roles of KMT2 writer enzymes in homeostatic scaling

The KMT2 H3K4 writers all regulate the same chromatin mark, raising the question of how each enzyme contributes uniquely to homeostatic scaling. One possibility is that the gene targets of each enzyme are non-overlapping. Indeed, distinct sets of genes were dysregulated when mouse *Kmt2a* and *Kmt2b* were ablated in adult excitatory neurons via expression of CaMKII Cre (Kerimoglu *et al.,* 2017). An alternative, though not mutually exclusive, possibility is that different KMT2 writers are recruited at different stages in the course of chronic activity changes. Consistent with this possibility, we find robust changes in global mono-methylation of H3K4 (H3K4me1) driven in the early phases of network hyperactivation, the vast majority of which appears to be mediated by KMT2A. Moreover, leveraging the KMT2A-specific inhibitor MM401, we found that KMT2A is required during this early period of hyperactivity (first 4 hrs) for the induction of synaptic downscaling, but is not required at later times. Thus, as is the case of KMT2A, specific KMT2 writers may be engaged at different times to mediate distinct aspects of coupling transcriptional outputs to synaptic adaptations. Our results further demonstrate that a major role of this early KMT2A activation is to sculpt the expression of late-response genes that are necessary for synaptic downscaling.

Our studies also allowed us to mechanistically dissect the induction phase of synaptic downscaling with its maintenance. We found that the induction of synaptic downscaling required the early activation of KMT2A but once established, downscaling maintenance was independent of KMT2A even over several days. Most studies have focused mechanistic analyses on the induction phase, and only a small number have examined how these homeostatic adaptations are maintained. Our results are consistent with other studies demonstrating that synaptic scaling persists across the time-course of extended activity shifts but is the first to demonstrate key mechanistic differences between the induction and maintenance phases of synaptic downscaling. This distinction is important, as recent work demonstrates that maintaining homeostatic plasticity can protect motoneurons against neurodegeneration in models of Amyotrophic Lateral Sclerosis (Orr *et al*., 2020). At present, it is not clear if the shift towards KMT2A-independence in synaptic downscaling reflects a general shift away from transcriptional dependence or if other gene expression regulatory mechanisms take over at that stage.

### KMT2A regulates transcription of activity-responsive genes during downscaling

The present work describes the role of KMT2A in neuronal activity-dependent transcription for the first time. Previous work found that KMT2A is required for full expression of the well-characterized IEG *Arc* in the mouse prefrontal cortex during a working memory task (Jakovcevski *et al*., 2015). However, in this study, Arc expression was reduced even before the training session, leaving it unclear whether KMT2A is required for experience-dependent Arc induction. Using genome-wide profiling of transcriptional dynamics, our Bru-seq data revealed a key role for the early activation of KMT2A in regulating a component of the transcriptional program induced by network hyperactivity. The most straight forward possibility is that early KMT2A activation underlies the transcription of early response genes, but we found that both ERGs and LRGs were regulated by KMT2A, with a significantly greater effect on LRGs. Two scenarios could underlie the higher sensitivity of LRGs to KMT2A blockade. First, KMT2A-sensitive LRGs could themselves be dependent on KMT2A-regulated ERGs. For example, these ERGs encode transcription factors such as Fos and NPAS4 that induce a second transcriptional wave (Yap and Greenberg, 2018). On the other hand, we did not find significant misregulation of Fos and NPAS4 when KMT2A was inhibited, though we did observe altered transcription of other transcription factor genes such as Ahrr and Hes5 as well as translational regulators including ribosomal genes (Table S1). Alternatively, deposition of H3K4me by KMT2A could regulate transcription of LRGs directly. Neuronal activity can drive long-lasting changes in chromatin accessibility (Fernandez-Albert *et al*., 2019), and early H3K4me placement via KMT2A might promote a priming effect on LRG transcription. Supporting this possibility, earlier work showed an interaction between KMT2A and CBP (Ernst *et al*., 2001), a key histone acetyltransferase for activity-dependent transcription via H3K27 acetylation. These scenarios are not mutually exclusive.

The relationship between KMT2 catalytic activity and gene transcription is complex. Many studies describe correlations between the loss of KMT2 enzymes, alterations in H3K4me, and changes in transcription, but these are not strong in general. In addition, H3K4me can be placed as a consequence of transcription (Hui Ng *et al*., 2003)(Soares *et al*., 2017). Many factors likely contribute to this complexity, including the impact of other chromatin regulatory marks, the presence of multiple H3K4 writers (and erasers), and the three distinct states of H3K4me itself (mono-, di-, or tri-methylation). Recent work shows that cell type is another variable for H3K4me placement by KMT2C and KMT2D (Xie *et al*., 2023). However, our results are consistent with studies showing instructive roles of H3K4me in transcriptional control after acute ablation of H3K4me writer enzymes or associated proteins using a drug-induced degron system (Hughes, Kelley and Klose, 2020)(Wang *et al*., 2023). We find that only a subset of hyperactivity-induced genes are KMT2A-dependent, which suggests that the combinatorial action of multiple H3K4me writers is likely important for tight control over the broader set of activity-responsive genes. Our Bru-seq data also identified an unexpected role for KMT2A in activity-dependent transcriptional suppression, again primarily directed towards LRGs. Previous work found that KMT2A can recruit histone deacetylases and polycomb machinery to negatively regulate transcription (Xia *et al*., 2003), so it is unclear if this suppression is a direct effect of KMT2A-dependent H3K4me deposition. Further investigation is warranted to determine how KMT2A interacts with other KMT2 writers and erasers to dynamically regulate H3K4me and shape activity-dependent transcriptional outputs.

How do the KMT2A-dependent genes contribute to synaptic downscaling? It is likely that a clearer answer to this question will emerge when the transcriptional control conferred by other KMT2 writers involved in downscaling (KMT2B, F, and G) is defined. But even an examination of just KMT2A-sensitive LRGs raises some interesting possibilities. These genes include *Pak1 and SORCS1*, which are both known to be important regulators of AMPAR trafficking (Hussain *et al*., 2015)(Ribeiro *et al*., 2019)(Savas *et al*., 2015). Furthermore, another KMT2A-dependent LRG is Sorting nexin 14 (SNX14), which interacts with the phosphoinositide PI3,5P2, generated by the lipid kinase, PIkfyve (Rivero-Ríos and Weisman, 2022), and has been shown to regulate neuronal excitability and synaptic function (Huang *et al*., 2014). We previously reported that shRNA-mediated knockdown or chemical inhibition of Pikfyve impairs long-term depression and homeostatic downscaling by disrupting AMPAR trafficking in hippocampal neurons (McCartney *et al*., 2014).

The emergence of six distinct H3K4 writer enzymes in mammals has remained somewhat of a mystery. These enzymes may have substantial functional redundancy in certain contexts, but the catalytic function of each has been subject to strong selection pressure, suggesting their unique signaling is functionally significant in other contexts. The clinical literature underscores this point – all six KMT2 writers are associated with neurodevelopmental disorders, suggesting that this unique function is likely neuronal. Our results define an important division of labor among H3K4me writers in homeostatic scaling that reveals the importance of combinatorial signaling in this enzyme family for coupling chronic changes in neural activity with synaptic adaptations via fine-tuning activity-dependent transcriptional control.

## Material and Methods

### Mouse

The ΔSET*Kmt2a* mouse line carries a deletion in the genome region encoding the catalytic SET domain (Terranova et al., 2006). *Kmt2a*^flox/flox^ (exons 8 and 9) was obtained as previously described (Jude et al., 2007). *Kmt2d* ^flox/flox^ mouse line carries a deletion in the catalytic SET domain which results in destabilization of the entire protein (Jang et al., 2019). All three mouse lines were back crossed with C57BL/6J (Jackson Laboratory). All animal use followed NIH guidelines and was in compliance with the University of Michigan Committee on Use and Care of Animals.

### Plasmids

pCAG-Cre:GFP was a gift from Dr. Connie Cepko (Addgene plasmid #13776). The retroviral pMSCVhyg plasmid expressing wild type (WT) histone H3.3 and Lys4-to-Met mutant expressing plasmids were gifts from Dr. Kai Ge (Addgene plasmid # 128740, # 128741). The following plasmids were used for the target-specific knockdown of the Kmt2 family. sh-*Kmt2a*: TRCN0000310900, sh-*Kmt2b*: TRCN0000034408, sh-*Kmt2c*#1: TRCN0000238937, sh-*Kmt2c*#2: TRCN0000238935, sh-*Kmt2d*: TRCN0000239233, sh-*Kmt2f*#1: TRCN0000225919, sh-*Kmt2f*#2: TRCN0000225920, sh-*Kmt2g*: TRCN0000095450 (Sigma MISSION). The non-targeting scramble sequence were used for a control (Sigma MISSION, sh-Ctrl: SHC202).

### Lentivirus preparation and transfection

Lentivirus was generated as previously described (Garay et al., 2020). Briefly, pLKO plasmids containing shRNA sequence to specifically target each KMT2s were co-transfected with psPAX2 and pMD2.G into HEK 293T cell line to generate lentivirus. The virus particles containing media were collected 24 h and 48 h after transfection and were concentrated using Lenti-X concentrator (Takara, 631232). The virus containing media was resuspended in Neurobasal media and tittered by puromycin selection using HEK 293T cell line. Transfection was performed as previously described (Garay et al., 2020). 1 µM of GFP and sh-RNA containing plasmids were co-transfected into primary cultured neurons with lipofectamine 2000 (Invitrogen), according to the manufacture protocol, at the indicated time described in the results.

### Primary culture

Primary hippocampal neurons were prepared as previously described (Henry et al., 2012). Briefly, hippocampi were dissected from postnatal (P0-3) CD1 mouse or Sprague-Dawley rat (Charles River) pups were dissociated and plated with growth media, including Neurobasal-A medium supplemented with B27 (Invitrogen), at a density of 230-460 mm^2^ in poly-D-lysine-coated glass-bottom petri dishes (Mattek) and cultured until used. Both sexes were mixed and used.

### Electrophysiology

Whole-cell patch-clamp recordings were performed as previously described (Garay et al., 2020). Briefly, whole-cell patch-clamp recordings were obtained from pyramidal neurons detected based on morphology. To record miniature excitatory postsynaptic current (mEPSC), micro-pipettes with 3-7 MΩ were filled with the following internal solution (in mM): 100 Cs-gluconate, 0.2 EGTA, 5 MgCl2, 2 MgATP, 0.3 GTP (pH 7.20). External solution includes the following (in mM): 119 NaCl, 5 KCl, 2 CaCl2, 2 MgCl2, 30 glucose, and 10 HEPES. For synaptic scaling, either 1 µM TTX or 40 µM bicuculline were treated 24-48 h before recording. Recordings were performed using Olympus IX-70 inverted microscopy, Multiclamp 700B (Axon Instruments), pCLAMP10 software (Axon Instruments), and Digidata 1440. Typically, recordings exceeding Ra < 25 MΩ or increasing 20% were excluded from further analysis. Minianalysis (Synaptosoft) and Clampfit 10.3 (Molecular Devices) were used for data analysis.

### Immunocytochemistry

Staining was conducted as previously described (Garay et al., 2020). Briefly, cells were fixed in 4% paraformaldehyde, permeabilized with 0.1% Triton X-100 in phosphate buffer saline (PBS), then blocked with 2 % bovine albumin serum for 1 h in room temperature. Primary antibodies for H3K4me1 (Abcam ab8895, 1:300), H3K4me3 (Abcam ab8580, 1:300), MAP2 (Sigma Aldrich, 1:5000), KMT2A (Cell signaling Technology 141975, 1:100), KMT2D (1:300) were incubated overnight at 4 °C. Alexa 555-Alexa 647-conjugated secondary antibodies were used for visualizing the staining. DAPI was used for staining the nucleus.

### Confocal microscopy

All imaging was conducted as previously described with minor modifications (Henry et al., 2012). In brief, inverted Olympus FV 1000 laser-scanning confocal microscope with a Plan-Apochromat 63 ×/1.4 oil objective was used (1-3 × digital zoom). Identical parameters were used through the same experiments. ImageJ were used to process the images. Images were analyzed on maximal intensity z-projections. Average integrated intensity was calculated in the nucleus. Generally, data were analyzed twice, including a trial performed by blinded undergraduate and high school students.

### Bromouridine sequencing (BrU-seq)

BrU-seq was conducted as previously described with modification (Garay et al., 2020). In brief, on DIV17, cortical cultures derived by CD1 mice were pretreated with either DMSO or 25 μM MM401. After 30min of preincubation with the reagents, cells were treated with bicuculline-methiodide (Abcam, ab120108, 40 μM) or vehicle (sterile water) for 1 h or 4 h. All cells were treated with 2 mM bromouridine (Bru, Sigma, 18670, dissolved in PBS) for the last 30 min of incubation. Cultures were lysat in Tri-reagent BD (Sigma, T3809) and frozen for future procedures. RNA was purified by phenol-chloroform extraction and isopropaonol precipitation, treated with DNase-I (NEB, M0303) then fragmented by high-magnesium, high-temperature incubation. One 1 g of Bru-containing RNA was used for library preparation. The library was prepared in collaboration with the BrU-seq lab of the University of Michigan (Paulsen et al., 2013, 2014). Finary, samples were sequenced by 150 bp paired-end on Illumina Novaseq. Experiments were conducted in three biological replicates with mixed sex.

### BrU-seq Analysis

First, the quality of the sequencing data was confirmed by FastQC. Then, reads were mapped to mm10 reference genome using Bowtie2 (Langmead and Salzberg 2012 et al., 2012) and annotated with Tophat2 (Kim et al., 2013). BrU-seq signals were quantified by FeatureCounts (Liao et al., 2014). Differentially expressed genes were identified using DESeq2 (Love et al., 2014). We also used DESeq2 to calculate the log2 fold change. Gene ontology enrichment pathways were obtained using ShinyGO 0.77. Typically, we used adjusted p-value (padj) < 0.05 for significance. Since, in general, the H3K4 methyl enzymes’ effect on transcription is not strong, we relaxed the threshold for detecting MM401-sensitive genes (padj < 0.03).

### Statistics

No statistical methods were used to predetermine sample size. Sample sizes were determined based on our previous work.

## Acknowledgements

We thank Cindy Carruthers and Christian Althaus for preparing neuronal cultures. We thank the members of the Iwase and Sutton labratories for their insights and suggestions. This work was supported by R01-NS129198 M.A.S., R01-NS097498 to M.A.S, and R21-NS125449, R21-MH127485 and R01-NS116008 to S.I., SMS research foundation to S.I. and M.A.S. T.T. was supported by a Parents and Researchers Interested in Smith-Magenis Syndrome (PRISMS) post-doctoral fellowship. A.C. was supported by a National Institutes of Health (NIH) National Research Service Award (NRSA) fellowship (18-PAF03228).

## Data availability

The Bru-seq data are available at NCBI Gene Expression Omnibus (<u>GSE239471</u>). The scripts used for Bru-seq analysis are available at https://github.com/tsukahar/DoL_project.

## Author Contribution

Conceptualization, T.T., S.I., and M.A.S.; Methodology, T.T., S.K., K.B., S.I., M.A.S.; Validation, T.T., S.K., B.L.S.; Formal Analysis, T.T., S.K., B.L.S., S.I.; Investigation, T.T., S.K., B.L.S.; Writing—Original Draft, T.T., S.I., and M.A.A.; Writing—Review & Editing: T.T., S.K., K.B., A.C., B.L.S., Y.D., S.I., and M.A.S.; Visualization, P.M.G., A.C., T.T., J.C.R.D., R.K., and M.A.W.; Supervision, T.T., S.I., and M.A.S.; Project Administration, T.T., S.I., and M.A.S.; Funding Acquisition, T.T., S.I., and M.A.S.

## Supplementary Figure Legend

**Figure S1.**
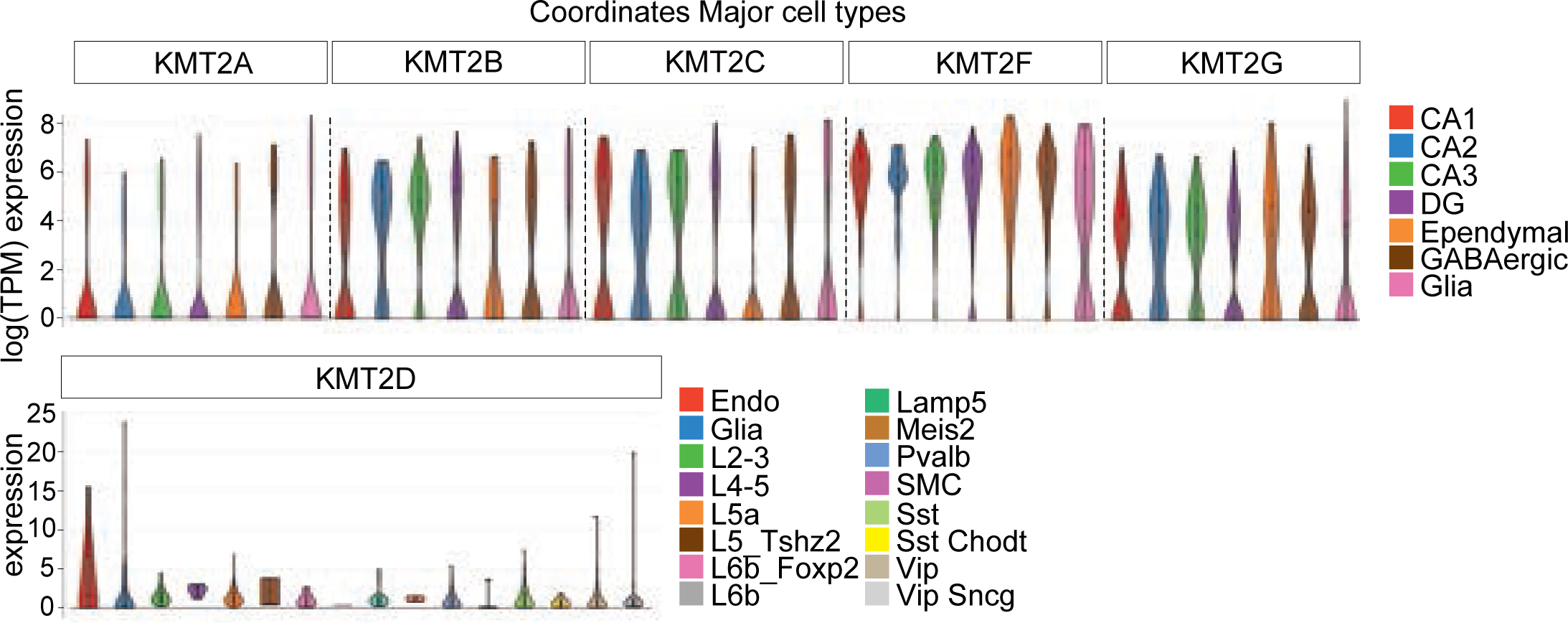
Single-cell RNA-seq reveals that all six members in the KMT2 family are ubiquitously expressed in most cell types in the brain. Violin plots display mean log (TPM) expression of the KMT2 family members in mouse hippocampal and cortical cells. Data was visualized using Single Cell Portal (Habib *et al*., 2016).

**Figure S2.**
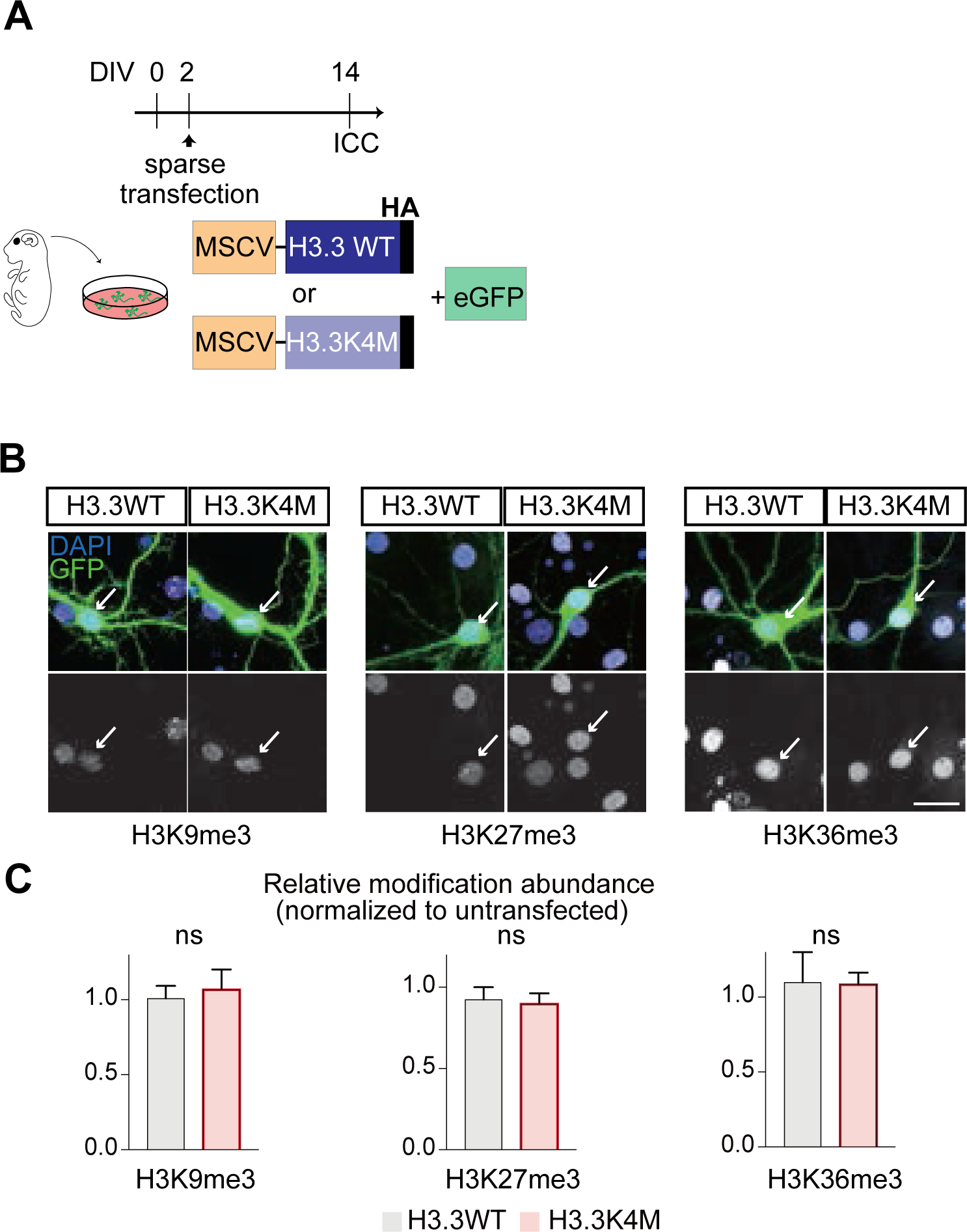
Ectopic expression of H3.3K4M doesn’t affect neighboring histone modifications. (A) Schema of experimental paradigm. (B) Representative images of H3K9me3 (white; left), H3K27me3 (white; middle), H3K36me3 (white; right) in H3.3WT or H3.3K4M transfected neurons. Scale bar 20 µm. (C) Bar graph of the relative modification abundance (normalized to untransfected neurons) of H3K9me3 (N=19-16), H3K27me27 (N=16-20), and H3K36me3 (N=17-17) in H3.3WT or H3.3K4M transfected neurons. White arrow labels transfected neurons. Mean ± S.E.M. Unpaired t-test. ns: p>0.05. (D) Representative miniature excitatory postsynaptic currents (mEPSC) traces from H3.3WT and H3.3K4M. Scale bar 125 ms, 20 pA. (F-H) Box plots of average mEPSC amplitude and the cumulative probability curve of mEPSC amplitude. mEPSC frequency (Hz) and decay time (ms). From here on, box plots indicate the 1st and 3rd quartiles with the median. Whiskers show the minimum and maximum data points.

**Figure S3.**
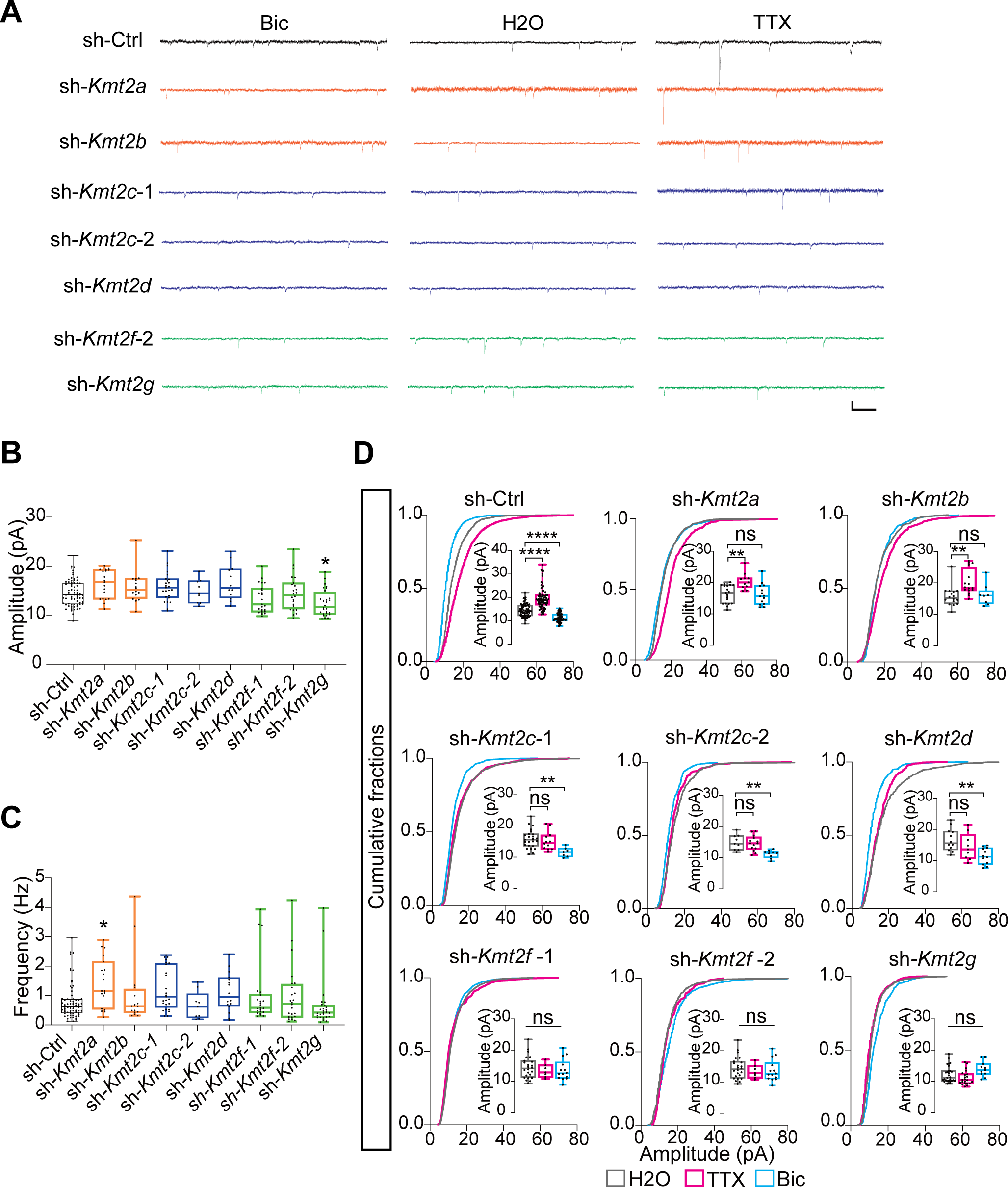
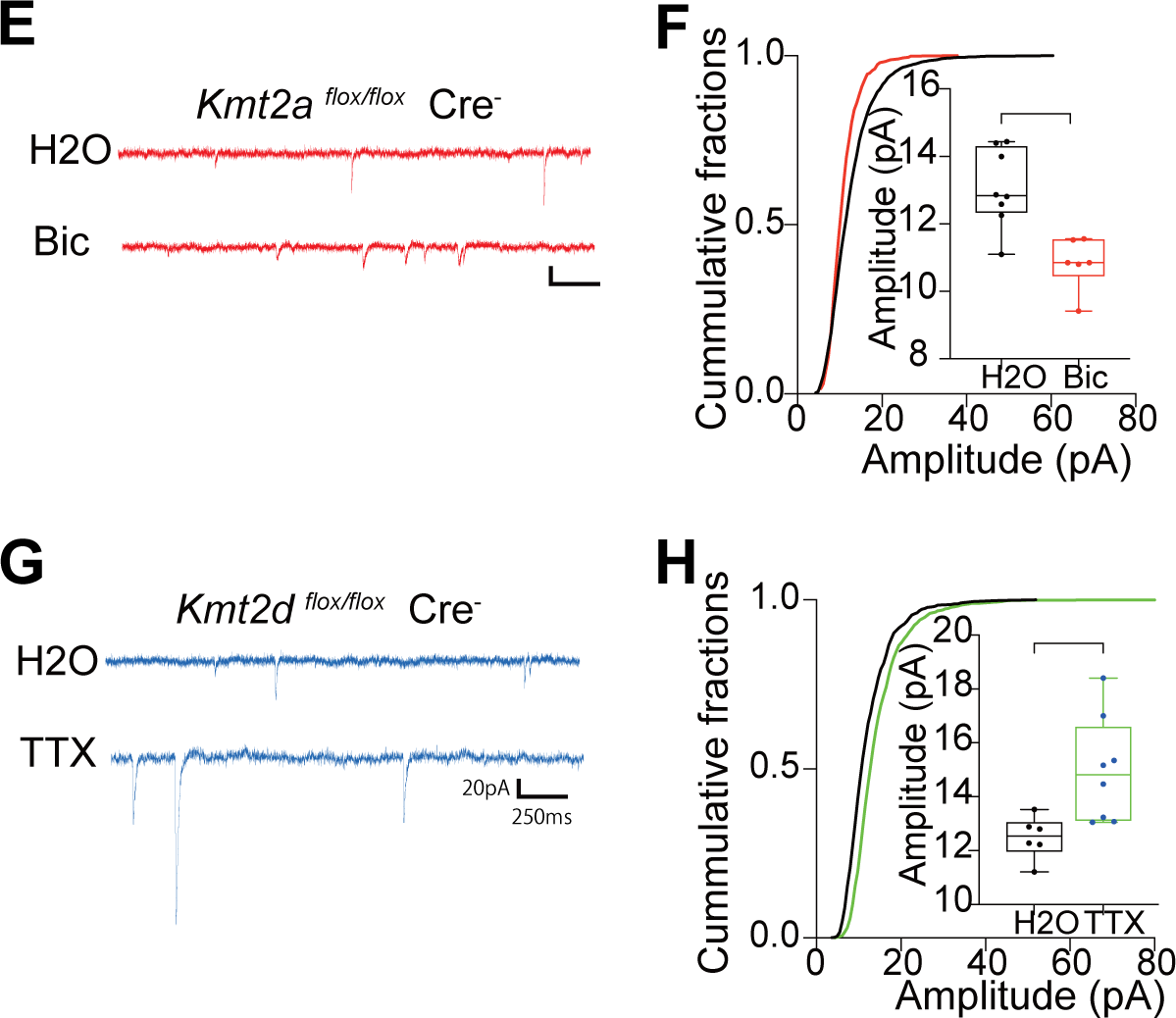
RNAi screening and conditional knock-out experiments unveil a division of labor among the KMT2 family members in the regulation of homeostatic synaptic scaling. (A) Representative traces for the mEPSC in neurons transfected with sh-RNAs that targets specific member in the KMT2 family. Hippocampal primary neurons were transfected with sh-RNAs at DIV 12 and further treated with H2O, TTX, or Bic to evoke synaptic scaling for 24∼48 h 2 days after transfection. (B-C) Box plots of average mEPSC amplitude and frequency (Hz) in neurons transfected with sh-RNAs targeting specific members in the KMT2 family. N=72,19,16,25,9,12,23,24,28 from the left. *Kmt2g*-KD exhibited moderate decrease in amplitude (P=0.040). *Kmt2a*-KD exhibited an increase in frequency (P=0.013). KD of most members in the KMT2 family had no effect in basal mEPSC amplitude and frequency. One-way ANOVA followed by post hoc Dunnett’s test. *p < 0.05. (D) Box plots of average mEPSC and cumulative probability curve of mEPSC amplitude in neurons transfected with sh-RNAs to KD individual KMT2s. Cultures were treated with either H2O, TTX, or Bic to induce synaptic scaling. (E-H) Cre negative neurons exhibit intact downscaling. (E) Representative mEPSC traces of *Kmt2a*^flox/flox^ Cre- neurons treated with either H2O or Bic. (F) Box plots of average mEPSC and cumulative probability curve of mEPSC amplitude in *Kmt2a*^flox/flox^ Cre- neurons (P = 0.0128). N=6-8. Unpaired t test. *P < 0.05 (G) Representative mEPSC traces of Kmt2a flox/flox Cre- neurons treated with either H2O or Bic. (H) Box plots of average mEPSC and cumulative probability curve of mEPSC amplitude in *Kmt2a*^flox/flox^ Cre- neurons. Unpaired t test. *P < 0.05.

**Figure S4.**
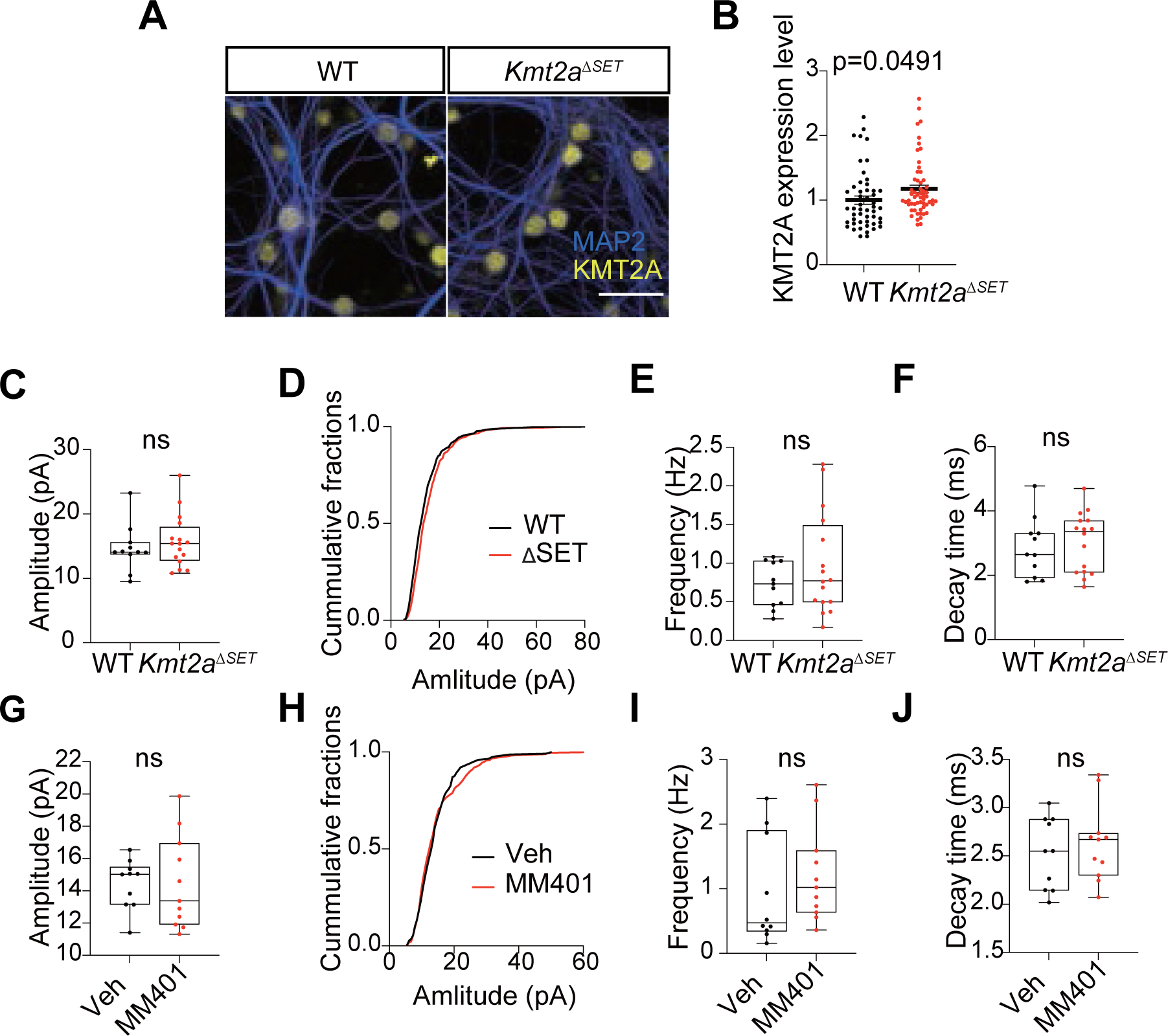
Genetic and pharmacological inhibition of the catalytic activity of KMT2A doesn’t affect basal glutamatergic transmission. (A) Representative images of WT and *Kmt2a*ΔSET mice derived hippocampal neurons immunolabeled with MAP2 (blue) and KMT2A (yellow). (B) Expression of KMT2A is not decreased in *Kmt2a*ΔSET mice neurons, compared to WT littermates. Mean ± S.E.M., P < 0.0491. Unpaired t-test. (C-F) Genetic deletion of the catalytic function of Kmt2a (*Kmt2a*ΔSET) did not exert any effects on mEPSC amplitude, frequency, and decay time in primary hippocampal neurons. (G-J) Acute inhibition of the catalytic function of KMT2A via the small molecule inhibitor MM401 did not exert any effects on basal mEPSC amplitude, frequency, and decay time in primary hippocampal neurons.

**Figure S5.**
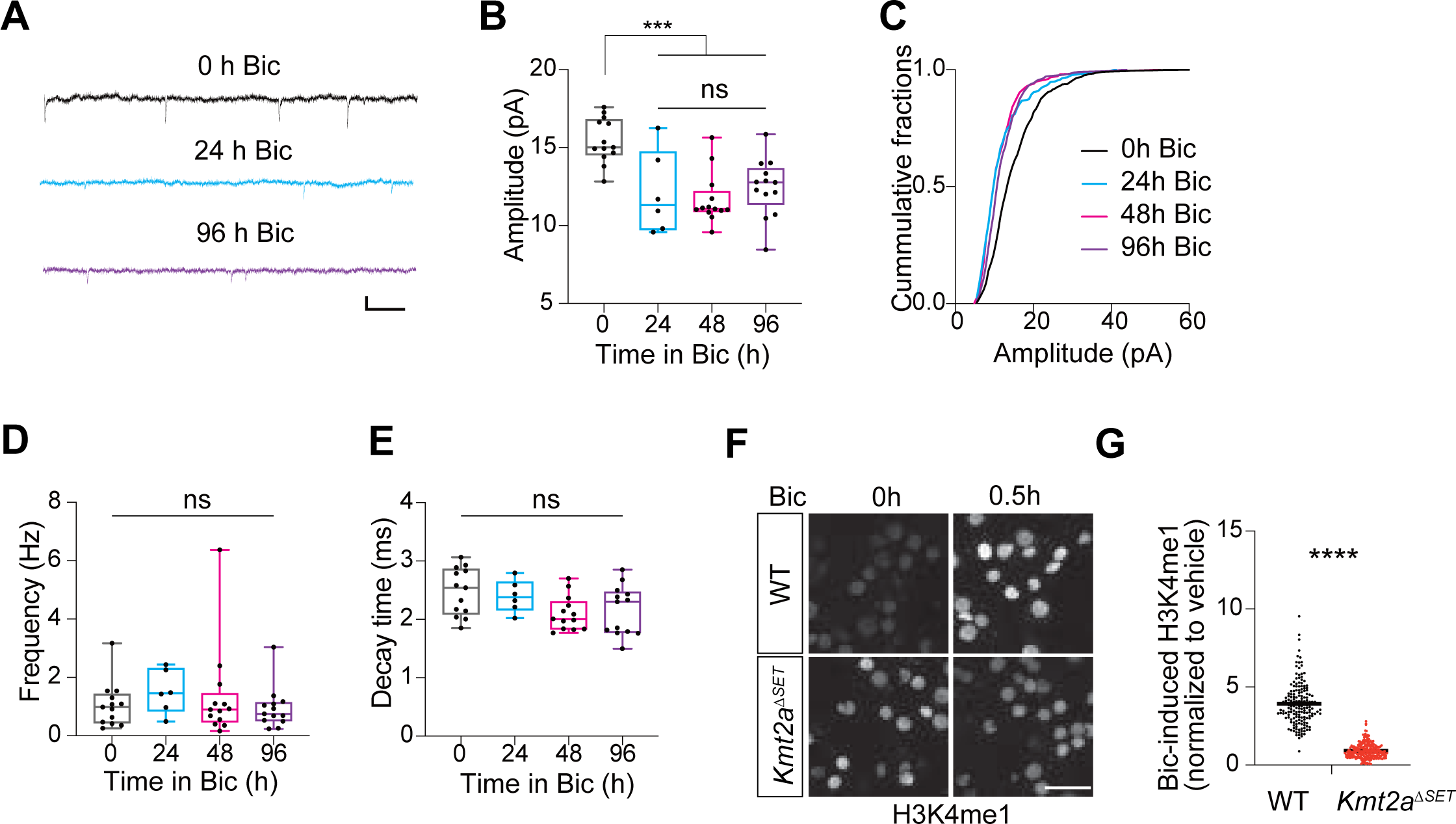
Maintenance of homeostatic downscaling exists. (A) Representative traces of mEPSC treated with 0 h, 24 h, and 96 h Bic. Scale bar 20 pA, 150 ms. (B and C) Box plot and cumulative probability curve of mEPSC amplitude comparing 0 h, 24 h, and 96 h Bic treatment. One-way ANOVA followed by Tukey’s multiple comparisons tests. ***P < 0.001. ns: P > 0.05. (D and E) Box plot of average mEPSC frequency comparing 0 h, 24 h, and 96 h Bic treatment. One-way ANOVA followed by Tukey’s multiple comparisons tests. ns: P > 0.05. (F) Representative images of hippocampal neurons treated with Bic for 0 h and 0.5 h. Neurons were derived from WT and *Kmt2a*ΔSET and labeled with anti-H3K4me1. (G) Average intensity of Bicuculline-induced (0.5 h Bic) H3K4me1 in WT and *Kmt2a*ΔSET. Mean ± S.E.M. ***P < 0.001. Unpaired t-test.

**Figure S6.**
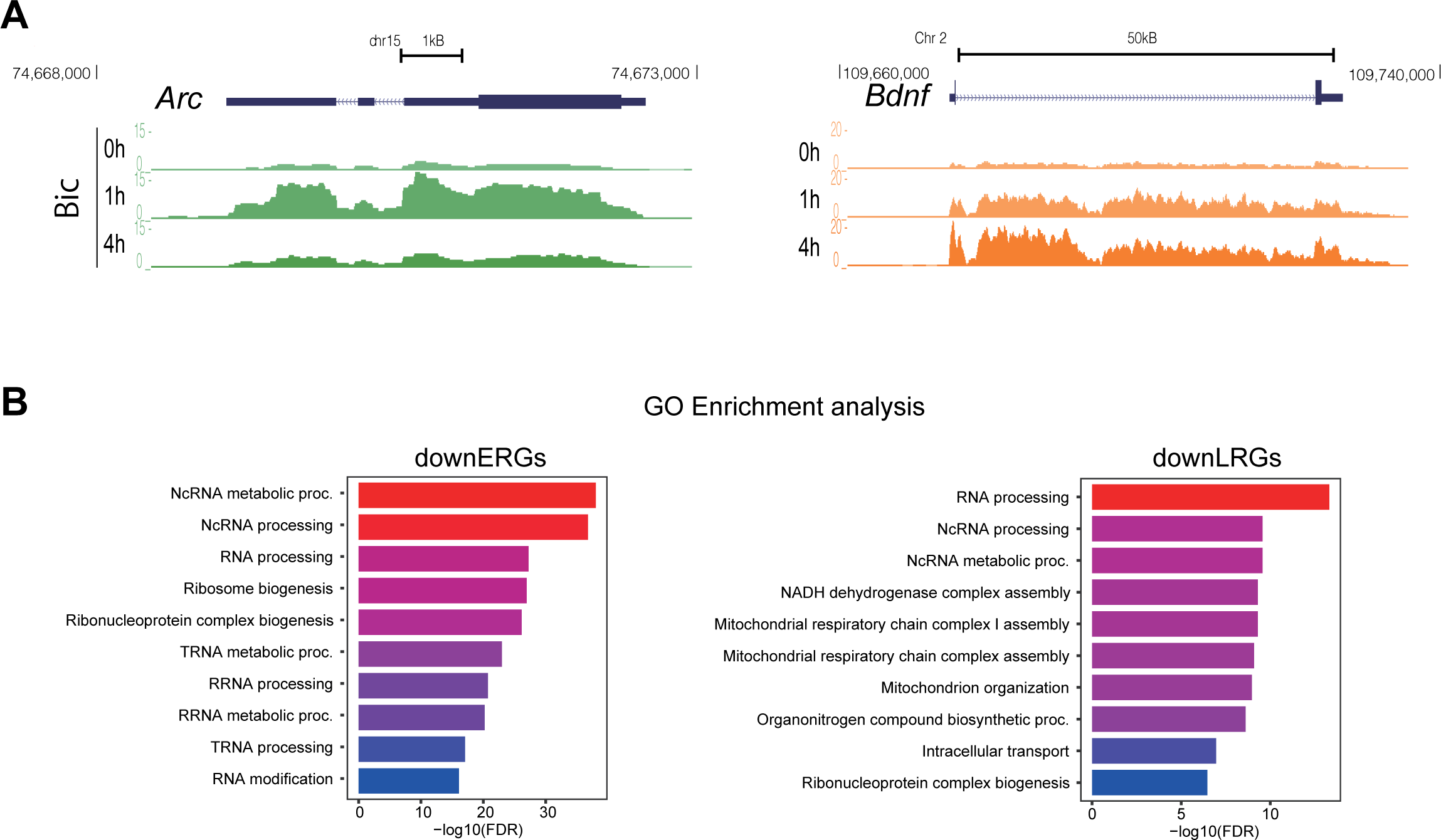
Nascent RNA-seq revealed that KMT2A inhibition attenuates the expression of late-responsive genes. (A) USCS genome browser shot of *Arc* and *Bdnf*, which represent up-regulated ERG (upERG) and LRG (upLRG), respectively. expression of *Arc* is transiently increased upon Bic treatment. On the other hand, *Bdnf* expression increased only after 4 h Bic treatment. The enriched intronic reads manifest that BrU-seq is capturing the nascent RNAs. (B) Gene-ontology enrichment analysis for downregulated ERGs (downERGs) and LRGs (downLRGs).

## References

Barski, A. et al. (2007) ‘High-Resolution Profiling of Histone Methylations in the Human Genome’, Cell, 129(4), pp. 823–837. doi: 10.1016/j.cell.2007.05.009.

Bateup, H. S. et al. (2013) ‘Excitatory/Inhibitory Synaptic Imbalance Leads to Hippocampal Hyperexcitability in Mouse Models of Tuberous Sclerosis’, Neuron. Elsevier, 78(3), pp. 510–522. doi: 10.1016/j.neuron.2013.03.017.

Benevento, M. et al. (2016) ‘Histone Methylation by the Kleefstra Syndrome Protein EHMT1 Mediates Homeostatic Synaptic Scaling’, Neuron. Elsevier Inc., 91(2), pp. 341–355. doi: 10.1016/j.neuron.2016.06.003.

Bledau, A. S. et al. (2014) ‘The H3K4 methyltransferase Setd1a is first required at the epiblast stage, whereas Setd1b becomes essential after gastrulation’, Development (Cambridge), 141(5), pp. 1022–1035. doi: 10.1242/dev.098152.

Cenik, B. K. and Shilatifard, A. (2021) ‘COMPASS and SWI/SNF complexes in development and disease’, Nature Reviews Genetics. Springer US, 22(1), pp. 38–58. doi: 10.1038/s41576-020-0278-0.

Chiu, S. L. et al. (2017) ‘GRASP1 Regulates Synaptic Plasticity and Learning through Endosomal Recycling of AMPA Receptors’, Neuron. Elsevier Inc., 93(6), pp. 1405–1419.e8. doi: 10.1016/j.neuron.2017.02.031.

Dorighi, K. M. et al. (2017) ‘Mll3 and Mll4 Facilitate Enhancer RNA Synthesis and Transcription from Promoters Independently of H3K4 Monomethylation’, Molecular Cell. Elsevier Inc., 66(4), pp. 568–576.e4. doi: 10.1016/j.molcel.2017.04.018.

Ernst, P. et al. (2001) ‘MLL and CREB Bind Cooperatively to the Nuclear Coactivator CREB-Binding Protein’, Molecular and Cellular Biology, 21(7), pp. 2249–2258. doi: 10.1128/mcb.21.7.2249-2258.2001.

Fernandez-Albert, J. et al. (2019) ‘Immediate and deferred epigenomic signatures of in vivo neuronal activation in mouse hippocampus’, Nature Neuroscience. Springer US, 22(10), pp. 1718–1730. doi: 10.1038/s41593-019-0476-2.

Fowler, T., Sen, R. and Roy, A. L. (2011) ‘Regulation of primary response genes’, Molecular Cell. Elsevier Inc., 44(3), pp. 348–360. doi: 10.1016/j.molcel.2011.09.014.

Froimchuk, E., Jang, Y. and Ge, K. (2017) ‘Histone H3 lysine 4 methyltransferase KMT2D’, Gene. Elsevier, 627(June), pp. 337–342. doi: 10.1016/j.gene.2017.06.056.

Garay, P. M. et al. (2020) ‘RAI1 Regulates Activity-Dependent Nascent Transcription and Synaptic Scaling’, Cell Reports, 32(6). doi: 10.1016/j.celrep.2020.108002.

Gehre, M. et al. (2020) ‘Lysine 4 of histone H3.3 is required for embryonic stem cell differentiation, histone enrichment at regulatory regions and transcription accuracy’, Nature Genetics. Springer US, 52(3), pp. 273–282. doi: 10.1038/s41588-020-0586-5.

Glaser, S. et al. (2006) ‘Multiple epigenetic maintenance factors implicated by the loss of MII2 in mouse development’, Development, 133(8), pp. 1423–1432. doi: 10.1242/dev.02302.

Gupta, S. et al. (2010) ‘Histone methylation regulates memory formation’, Journal of Neuroscience, 30(10), pp. 3589–3599. doi: 10.1523/JNEUROSCI.3732-09.2010.

Habib, N. et al. (2016) ‘Div-Seq: Single-nucleus RNA-Seq reveals dynamics of rare adult newborn neurons’, Science, 353(6302), pp. 925–928. doi: 10.1126/science.aad7038.

Heintzman, N. D. et al. (2007) ‘Distinct and predictive chromatin signatures of transcriptional promoters and enhancers in the human genome’, Nature Genetics, 39(3), pp. 311–318. doi: 10.1038/ng1966.

Herz, H. M. et al. (2012) ‘Enhancer-associated H3K4 monomethylation by trithorax-related, the drosophila homolog of mammalian MLL3/MLL4’, Genes and Development, 26(23), pp. 2604–2620. doi: 10.1101/gad.201327.112.

Herz, H. M. et al. (2014) ‘Histone H3 lysine-to-methionine mutants as a paradigm to study chromatin signaling’, Science, 345(6200), pp. 1065–1070. doi: 10.1126/science.1255104.

Hiraide, T. et al. (2018) ‘De novo variants in SETD1B are associated with intellectual disability, epilepsy and autism’, Human Genetics. Springer Berlin Heidelberg, 137(1), pp. 95–104. doi: 10.1007/s00439-017-1863-y.

Hödl, M. and Basler, K. (2012) ‘Transcription in the absence of histone H3.2 and H3K4 methylation’, Current Biology, 22(23), pp. 2253–2257. doi: 10.1016/j.cub.2012.10.008.

Horvath, P. M. et al. (2021) ‘Report A subthreshold synaptic mechanism regulating BDNF expression and resting synaptic strength ll ll A subthreshold synaptic mechanism regulating BDNF expression and resting synaptic strength’, CellReports. ElsevierCompany., 36(5), p. 109467. doi: 10.1016/j.celrep.2021.109467.

Hu, D. et al. (2013) ‘The MLL3/MLL4 Branches of the COMPASS Family Function as Major Histone H3K4 Monomethylases at Enhancers’, Molecular and Cellular Biology, 33(23), pp. 4745–4754. doi: 10.1128/mcb.01181-13.

Huang, H. S. et al. (2014) ‘Snx14 regulates neuronal excitability, promotes synaptic transmission, and is imprinted in the brain of mice’, PLoS ONE, 9(5), pp. 1–9. doi: 10.1371/journal.pone.0098383.

Hughes, A. L., Kelley, J. R. and Klose, R. J. (2020) ‘Understanding the interplay between CpG island-associated gene promoters and H3K4 methylation’, Biochimica et Biophysica Acta - Gene Regulatory Mechanisms. Elsevier, 1863(8), p. 194567. doi: 10.1016/j.bbagrm.2020.194567.

Hui Ng, H., et al. (2003) ‘Targeted Recruitment of Set1 Histone Methylase by Elongating Pol II Provides a Localized Mark and Memory of Recent Transcriptional Activity genome-wide fashion (Kuo et al that is linked to subsequent events of mRNA production including capping, splicing’, Orphanides and Reinberg, 11, pp. 709–719.

Hussain, N. K. et al. (2015) ‘Regulation of AMPA receptor subunit GluA1 surface expression by PAK3 phosphorylation’, Proceedings of the National Academy of Sciences of the United States of America, 112(43), pp. E5883–E5890. doi: 10.1073/pnas.1518382112.

Jakovcevski, M. et al. (2015) ‘Neuronal Kmt2a/Mll1 histone methyltransferase is essential for prefrontal synaptic plasticity and working memory’, Journal of Neuroscience, 35(13), pp. 5097–5108. doi: 10.1523/JNEUROSCI.3004-14.2015.

Jang, Y. et al. (2019) ‘H3.3K4M destabilizes enhancer H3K4 methyltransferases MLL3/MLL4 and impairs adipose tissue development’, Nucleic Acids Research. Oxford University Press, 47(2), pp. 607–620. doi: 10.1093/nar/gky982.

Jones, W. D. et al. (2012) ‘De novo mutations in MLL cause Wiedemann-Steiner syndrome’, American Journal of Human Genetics, 91(2), pp. 358–364. doi: 10.1016/j.ajhg.2012.06.008.

Koemans, T. S. et al. (2017) ‘Functional convergence of histone methyltransferases EHMT1 and KMT2C involved in intellectual disability and autism spectrum disorder’, PLoS Genetics, 9(1), pp. 1–24. doi: 10.1371/journal.pbio.1000569.

Lee, J. E. et al. (2013) ‘H3K4 mono-And di-methyltransferase MLL4 is required for enhancer activation during cell differentiation’, eLife, 2013(2), pp. 1–25. doi: 10.7554/eLife.01503.

Li, D. D. et al. (2016) ‘High-affinity small molecular blockers of mixed lineage leukemia 1 (MLL1)-WDR5 interaction inhibit MLL1 complex H3K4 methyltransferase activity’, European Journal of Medicinal Chemistry. Elsevier Masson SAS, 124, pp. 480–489. doi: 10.1016/j.ejmech.2016.08.036.

Mao, W. et al. (2018) ‘Activity-Induced Regulation of Synaptic Strength through the Chromatin Reader L3mbtl1’, Cell Reports. ElsevierCompany., 23(11), pp. 3209–3222. doi: 10.1016/j.celrep.2018.05.028.

McCartney, A. J. et al. (2014) ‘Activity-dependent PI(3,5)P2 synthesis controls AMPA receptor trafficking during synaptic depression’, Proceedings of the National Academy of Sciences of the United States of America, 111(45), pp. E4896–E4905. doi: 10.1073/pnas.1411117111.

Mishra, B. P. et al. (2014) ‘The histone methyltransferase activity of MLL1 is dispensable for hematopoiesis and leukemogenesis’, Cell Reports. The Authors, 7(4), pp. 1239–1247. doi: 10.1016/j.celrep.2014.04.015.

Morgan, M. A. J. and Shilatifard, A. (2020) ‘modifications in transcriptional regulation’, Nature Genetics. Springer US, 52(December). doi: 10.1038/s41588-020-00736-4.

Ng, S. B. et al. (2010) ‘Exome sequencing identifies MLL2 mutations as a cause of Kabuki syndrome’, Nature Genetics, 42(9), pp. 790–793. doi: 10.1038/ng.646.

Orr, B. O. et al. (2020) ‘Presynaptic Homeostasis Opposes Disease Progression in Mouse Models of ALS-Like Degeneration: Evidence for Homeostatic Neuroprotection’, Neuron. Elsevier Inc., pp. 1–17. doi: 10.1016/j.neuron.2020.04.009.

Paulsen, M. T. et al. (2013) ‘Coordinated regulation of synthesis and stability of RNA during the acute TNF-induced proinflammatory response’, Proceedings of the National Academy of Sciences of the United States of America, 110(6), pp. 2240–2245. doi: 10.1073/pnas.1219192110.

Paulsen, M. T. et al. (2014) ‘Use of Bru-Seq and BruChase-Seq for genome-wide assessment of the synthesis and stability of RNA’, Methods. Elsevier Inc., 67(1), pp. 45–54. doi: 10.1016/j.ymeth.2013.08.015.

Rao, R. C. and Dou, Y. (2015) ‘Hijacked in cancer: The KMT2 (MLL) family of methyltransferases’, Nature Reviews Cancer. Nature Publishing Group, 15(6), pp. 334–346. doi: 10.1038/nrc3929.

Ribeiro, L. F. et al. (2019) SorCS1-mediated sorting in dendrites maintains neurexin axonal surface polarization required for synaptic function, PLoS Biology. doi: 10.1371/journal.pbio.3000466.

Rickels, R. et al. (2017) ‘Histone H3K4 monomethylation catalyzed by Trr and mammalian COMPASS-like proteins at enhancers is dispensable for development and viability’, Nature Genetics. Nature Publishing Group, 49(11), pp. 1647–1653. doi: 10.1038/ng.3965.

Rivero-Ríos, P. and Weisman, L. S. (2022) ‘Roles of PIKfyve in multiple cellular pathways’, Current Opinion in Cell Biology. Elsevier Ltd, 76(5), p. 102086. doi: 10.1016/j.ceb.2022.102086.

Savas, J. N. et al. (2015) ‘The Sorting Receptor SorCS1 Regulates Trafficking of Neurexin and AMPA Receptors’, Neuron, 87(4), pp. 764–780. doi: 10.1016/j.neuron.2015.08.007.

Sedkov, Y. et al. (1999) ‘Molecular genetic analysis of the Drosophila trithorax-related gene which encodes a novel SET domain protein’, Mechanisms of Development, 82(1–2), pp. 171–179. doi: 10.1016/S0925-4773(98)00246-9.

Soares, L. M. et al. (2017) ‘Determinants of Histone H3K4 Methylation Patterns’, Molecular Cell. Elsevier Inc., 68(4), pp. 773–785.e6. doi: 10.1016/j.molcel.2017.10.013.

Soden, M. E. and Chen, L. (2010) ‘Fragile X protein FMRP is required for homeostatic plasticity and regulation of synaptic strength by retinoic acid’, Journal of Neuroscience, 30(50), pp. 16910–16921. doi: 10.1523/JNEUROSCI.3660-10.2010.

Takata, A. et al. (2014) ‘Loss-of-Function Variants in Schizophrenia Risk and SETD1A as a Candidate Susceptibility Gene’, Neuron. Elsevier Inc., 82(4), pp. 773–780. doi: 10.1016/j.neuron.2014.04.043.

Terranova, R. et al. (2006) ‘Histone and DNA methylation defects at Hox genes in mice expressing a SET domain-truncated form of Mll’, Proceedings of the National Academy of Sciences of the United States of America, 103(17), pp. 6629–6634. doi: 10.1073/pnas.0507425103.

Turrigiano, G. (2012) ‘Homeostatic synaptic plasticity: Local and global mechanisms for stabilizing neuronal function’, Cold Spring Harbor Perspectives in Biology, 4(1), pp. 1–18. doi: 10.1101/cshperspect.a005736.

Turrigiano, G. G. et al. (1998) ‘Activity-dependent scaling of quantal amplitude in neocortical neurons’, Nature, 391(6670), pp. 892–896. doi: 10.1038/36103.

Tusi, B. K. et al. (2015) ‘Setd1a regulates progenitor B-cell-to-precursor B-cell development through histone H3 lysine 4 trimethylation and Ig heavy-chain rearrangement’, FASEB Journal, 29(4), pp. 1505–1515. doi: 10.1096/fj.14-263061.

Wang, C. et al. (2016) ‘Enhancer priming by H3K4 methyltransferase MLL4 controls cell fate transition’, Proceedings of the National Academy of Sciences of the United States of America, 113(42), pp. 11871–11876. doi: 10.1073/pnas.1606857113.

Wang, H. et al. (2023) ‘H3K4me3 regulates RNA polymerase II promoter-proximal pause-release’, Nature. Springer US, 615(7951), pp. 339–348. doi: 10.1038/s41586-023-05780-8.

Xia, Z. B. et al. (2003) ‘MLL repression domain interacts with histone deacetylases, the polycomb group proteins HPC2 and BMI-1, and the corepressor C-terminal-binding protein’, Proceedings of the National Academy of Sciences of the United States of America, 100(14), pp. 8342–8347. doi: 10.1073/pnas.1436338100.

Xie, G. et al. (2023) ‘MLL3/MLL4 methyltransferase activities control early embryonic development and embryonic stem cell differentiation in a lineage-selective manner’, Nature Genetics. Springer US, 55(4), pp. 693–705. doi: 10.1038/s41588-023-01356-4.

Yap, E. L. and Greenberg, M. E. (2018) ‘Activity-Regulated Transcription: Bridging the Gap between Neural Activity and Behavior’, Neuron. Elsevier Inc., 100(2), pp. 330–348. doi: 10.1016/j.neuron.2018.10.013.

Yu, B. D. et al. (1995) ‘Altered Hox expression and segmental identity in Mll-mutant mice’, Nature, 378(November), pp. 505–508.

Yu, B. D. et al. (1998) ‘MLL, a mammalian trithorax-group gene, functions as a transcriptional maintenance factor in morphogenesis’, Proceedings of the National Academy of Sciences of the United States of America, 95(18), pp. 10632–10636. doi: 10.1073/pnas.95.18.10632.

Yu, H. et al. (2015) ‘Tet3 regulates synaptic transmission and homeostatic plasticity via DNA oxidation and repair’, Nature Neuroscience, 18(6), pp. 836–843. doi: 10.1038/nn.4008.

Yu, X. et al. (2019) ‘De Novo and Inherited SETD1A Variants in Early-onset Epilepsy’, Neuroscience Bulletin. Springer Singapore, 35(6), pp. 1045–1057. doi: 10.1007/s12264-019-00400-w.

Zech, M. et al. (2016) ‘Haploinsufficiency of KMT2B, Encoding the Lysine-Specific Histone Methyltransferase 2B, Results in Early-Onset Generalized Dystonia’, American Journal of Human Genetics. American Society of Human Genetics, 99(6), pp. 1377–1387. doi: 10.1016/j.ajhg.2016.10.010.

Zhong, X., Li, H. and Chang, Q. (2012) ‘MeCP2 phosphorylation is required for modulating synaptic scaling through mGluR5’, Journal of Neuroscience, 32(37), pp. 12841–12847. doi: 10.1523/JNEUROSCI.2784-12.2012.

